# Protein UFMylation regulates early events of ribosomal DNA double-stranded break response

**DOI:** 10.1101/2024.02.14.580383

**Authors:** Pudchalaluck Panichnantakul, Lisbeth C. Aguilar, Evan Daynard, Mackenzie Guest, Colten Peters, Jackie Vogel, Marlene Oeffinger

**Affiliations:** Institut de recherches cliniques de Montréal, Center for Genetic and Neurological Diseases, 110 avenue des Pins Ouest, Montréal, Québec, H2W 1R7; Division of Experimental Medicine, Faculty of Medicine, McGill University, Montréal, Québec, H4A 3J1; Department of Biology, Faculty of Medicine, McGill University, Montréal, Québec, H3A 1B1; Département de biochimie et médicine moléculaire, Faculté de Médicine, Université de Montréal, Québec, H3C 3J7

## Abstract

The highly repetitive and transcriptionally active ribosomal DNA (rDNA) genes are exceedingly susceptible to genotoxic stress. Induction of DNA double-strand breaks (DSBs) in rDNA repeats is associated with ATM-dependent rDNA silencing and nucleolar reorganization where rDNA is segregated into nucleolar caps. However, the regulatory events underlying this response remain elusive. Here, we identify protein UFMylation as essential for rDNA damage response in human cells. We further show the only UFM1-E3-ligase UFL1 and its binding partner DDRGK1 localize to nucleolar caps upon rDNA damage, and that UFL1 loss impairs ATM activation and rDNA transcriptional silencing, leading to reduced rDNA segregation. A first-ever analysis of nuclear and nucleolar UFMylation targets in response to DSBs induction further identified key DNA repair factors including ATM, in addition to chromatin and actin network regulators. Taken together, our data provides the first evidence of an essential role for UFMylation in orchestrating rDNA DSB repair.

## INTRODUCTION

The nucleolus is a dynamic phase-separated compartment within the nucleus which harbours ribosomal DNA (rDNA) genes, ribosomal RNA (rRNA), and hundreds of proteins^1–3^. In humans, rDNA genes form nucleolar organizing regions on the five acrocentric chromosomes in tandem repeats of ∼300 copies per cell^1,4^. Its high transcription frequency and repetitive sequences make rDNA exceedingly susceptible to genotoxic stress and double-stranded breaks (DSBs)^5,6^, and its instability has been attributed to recombination events, resulting in copy number variation that are observed in many human cancers where the natural variability in rDNA repeat numbers is increased by ∼20–80%^7–9^.

Unlike in the nucleoplasm, genotoxic ribosomal DNA stress leads to significant changes in nucleolar structure and the rDNA locus, including transcriptional repression via constitutive rDNA silencing through histone modifications and chromatin remodelling, and the segregation of damaged rDNA into nucleolar caps – crescent-shaped structures at the nucleolar periphery^10–12^. The segregation of damaged rDNA into nucleolar caps has been suggested to serve as prevention of inter-chromosomal recombination and, furthermore, to facilitate ribosomal DNA repair whereby nucleolar caps function as hubs for the recruitment of DNA repair proteins, away from nucleolar subregions where ribosomal RNA processing and assembly takes place^13^.

Several high-resolution microscopy studies have previously shown that rDNA DSB-induced repression of RNA Pol I transcription is dependent on the DNA repair serine/threonine kinases Ataxia-Telangiesctasia-Mutated (ATM) and Ataxia-Telangiesctasia-and-Rad3 related (ATR) as well as the downstream effector <u>hu</u>man <u>s</u>ilencing <u>h</u>ub (HUSH) complex, the latter of which was demonstrated to mediate H3K9Me3 deposition onto damaged rDNA, resulting in transcriptional silencing^10–13^. In addition, the nucleolar protein TCOF1 (treacle ribosome biogenesis factor 1, Treacle) has emerged as a critical factor for nucleolar cap formation in response to rDNA damage^12,14^. TCOF1 is essential for nucleolar cap formation in response to both RNA Polymerase (Pol) I inhibition and rDNA DSBs as TCOF1-depleted cells are unable to form nucleolar caps or silence rDNA transcription, while displaying reduced viability, increased apoptosis, and overall higher levels of genomic instability^12,14^. Upon rDNA DSB, TCOF1 furthermore facilitates the recruitment of the key DNA repair factors TOPBP1, and the <u>M</u>RE11-<u>R</u>AD50-<u>N</u>BS1 (MRN) complex to nucleolar caps, which is no longer observed in the absence of TCOF1^12,14–16^.

Orchestration of the early steps of the DNA damage response is believed to be regulated via posttranslational modifications, in particular phosphorylation events. In nuclear DNA DSB repair, ATM phosphorylates histone γH2AX at break sites, enabling the binding of MDC1 to damaged DNA^17,18^; this in turn leads to recruitment of NBS1 and activation of a positive feedback loop resulting in increased ATM activation and expansion of γH2AX phosphorylation around the DSB^17–20^. In contrast, upon generation of DSBs in rDNA genes, phosphorylation of TCOF1 by ATM at Ser1199 and Casein Kinase 2 at Thr210 facilitates recruitment of NBS1 and the MRN complex to nucleoli^12,21–23^. This recruitment subsequently promotes increasing levels of ATM activation in a positive feedback loop similar to that observed on nuclear chromatin^12,14,24^. After ATM activation, TCOF1 recruits TOPBP1 to the nucleolus, followed by ATR. These activation and recruitment steps are suggested to be required for complete rDNA transcriptional silencing and nucleolar segregation into caps to occur^12^.

Besides phosphorylation, other post-ranslational modification events were also shown to be essential for the regulation of genome maintenance pathways. For example, ubiquitination of ATMIN promotes its dissociation from ATM, which is a requirement for ATM interaction with NBS1 in response to ionizing radiation^25,26^. Moreover, in *Saccharomyces cerevisiae*, where damaged rDNA repeats must be moved to the nucleoplasm to enable repair by homologous recombination, release of rDNA from the nucleolus requires SUMOylation of the CLIP-cohibin tethering complex^27^.

More recent work has also shown UFMylation, the covalent conjugation of the ubiquitin-fold modifier 1 (UFM1) protein to lysine residues on target proteins via an enzymatic process involving the E1-like enzyme UBA5, the E2-like UFC1, and the E3-like UFL1^28–33^, to be implicated in nuclear DNA maintenance^34–37^. UFMylation of MRE11 was suggested to potentially promote MRN complex formation and its recruitment to genomic loci for their maintenance, such as telomeres^35^, while UFMylation of both MRE11 and histone H4 upon ionizing radiation-induced DSBs was suggested to facilitate ATM activation through a positive feedback loop involving SUV39H1/2 recruitment to nuclear DSBs, H3K9-trimethylation, and subsequent ATM signal amplification^34,36,37^. In that context, an MRN complex-dependent recruitment of the E3-ligase UFL1 to DNA damage foci has also been observed^34^. It is currently not known whether UFMylation is involved in ribosomal DNA damage.

Here, we use a genetic loss-of-function screen to determine genes required for cell viability upon ribosomal DNA damage and identify all components of the protein UFMylation pathway as essential for the ribosomal DNA damage response to double strand breaks in human cells. Our interactome and microscopy data further demonstrate that the UFM1-E3 ligase UFL1 and its binding partner DDRGK1 localize from the nucleoplasm into nucleolar caps, where they associate with TCOF1 upon induction of rDNA DSBs. Moreover, loss of the only known UFM1 E3-ligase, UFL1, leads to impaired ATM activation, decreased efficiency in rDNA transcriptional silencing, and reduced segregation of rDNA into nucleolar caps, in addition to reduced nucleolar levels of key DNA repair factors upon rDNA damage, suggesting that UFMylation is essential for a robust rDNA damage response, specifically its early steps. Furthermore, a first ever global analysis of nuclear and nucleolar UFMylation targets identified key components of the early rDNA damage response as well as several chromatin and cytoskeleton/actin network regulators as targets of UFMylation in response to DNA DSB induction. Taken together, our study provides the first evidence of an essential role for UFMylation in the regulation of early steps of rDNA DSB repair and identifies key nuclear UFMylation targets in response to rDNA and DNA DSB induction.

## RESULTS

### Protein UFMylation, H3.3, deposition, and the HUSH complex are essential for rDNA DSB repair

Although several studies have provided important insights into the cellular pathways involved in rDNA DSB repair, to date, we still have only a partial picture of how rDNA is repaired, in part because the full repertoire of factors involved in the repair process has yet to be elucidated^5,10–13,27,38–40^.

We therefore took a functional genomics approach to unbiasedly identify genes that are essential for rDNA DSBs repair by inducing a defined DSB within the rDNA using the fungal homing endonuclease I-PpoI which targets a 15-bp canonical site in all ∼300 copies of the 28S rDNA coding region and which has been extensively used to study rDNA damage ^5,10,12,41^. To control and track I-PpoI activity *in vivo*, we stably expressed an engineered HA-tagged I-PpoI cDNA fused to a destabilization domain (DD) and a 4-hydroxy-tamoxifen (4OHT)-inducible estrogen receptor (ER)-nuclear translocation domain, enabling controlled protein expression and nuclear localization upon addition of the ligands Shield-1 and 4-hydroxytamoxifen (4-OHT) (Figure S1A)^42^, in hTERT-RPE1 *p53-/-* Cas9 cells. Integration of this engineered I-Ppol into hTERT-RPE1 *p53-/-* Cas9 cells did not cause an increase in basal γH2AX protein levels, indicating that nuclear stress response remained at basal levels (Figure S1B-C), while stabilization and nuclear import of I-PpoI upon addition of Shield-1+4-OHT resulted in rDNA DSBs and nuclear cap formation in more than 80% of cells compared to less than 5% in DMSO-treated control cells as determined by immunofluorescent microscopy of TCOF1 (Figure 1A, B). These results furthermore phenocopied the nucleolar segregation observed upon transfection of hTERT-RPE1 *p53-/-* Cas9 cells with single guide (sg) RNAs targeting the 5’external transcribed spacer (5’ETS) region of the rDNA (Figure S1D)^41^.

**Figure 1.**
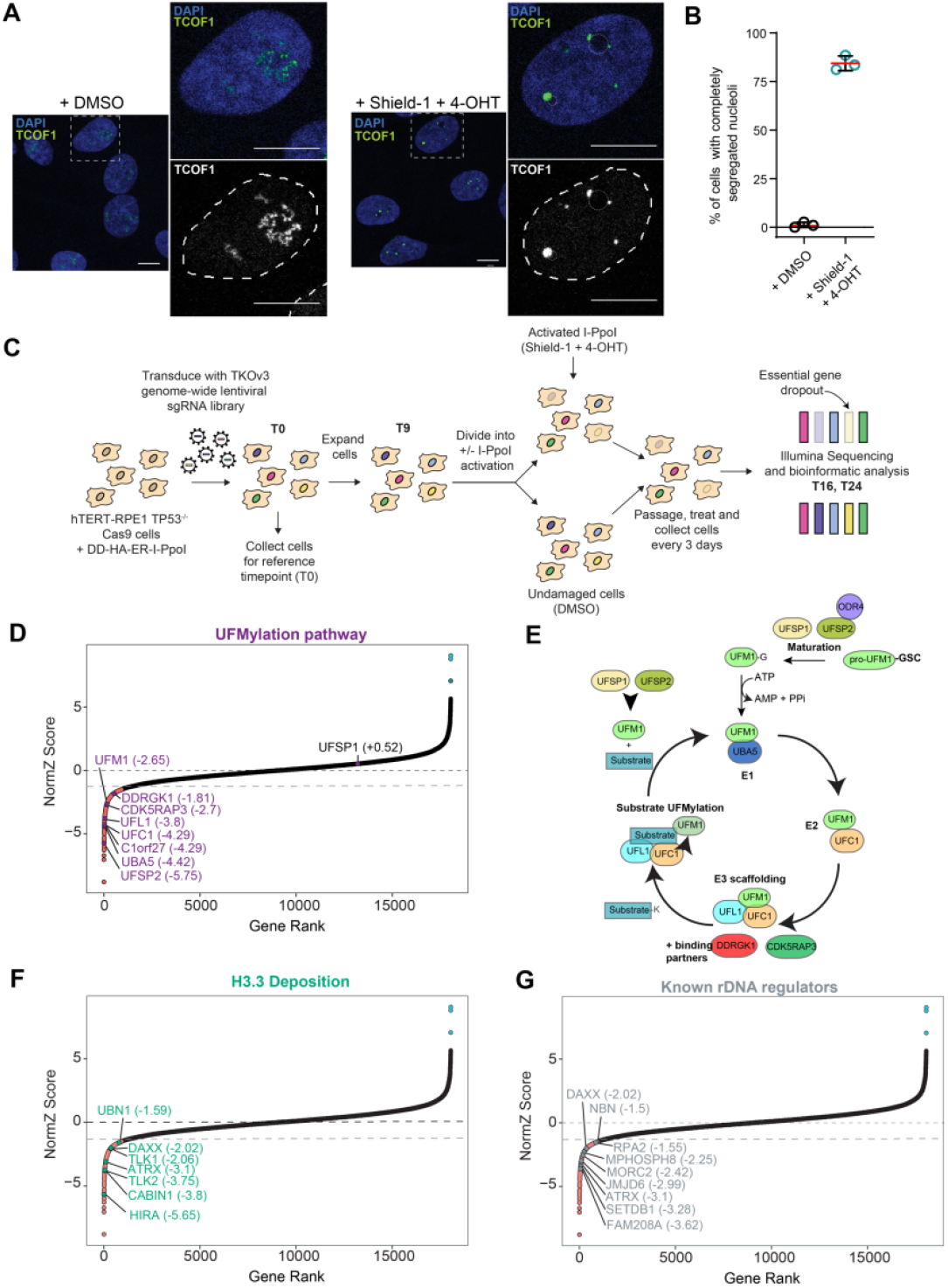
Genome-wide CRISPR-Cas9 screen identifies the protein UFMylation pathway, chromatin remodelers, and HUSH complex as essential to rDNA DSBs repair. **(A)** Immunofluorescent microscopy of hTERT-RPE1 *p53*^-/-^ Cas9 cells stably expressing DD-HA-ER-I-PpoI + DMSO or Shield-1+4-OHT for 6 hr. Dashed boxes show zoomed inlet. Green, endogenous TCOF1; DAPI, nucleus outlined with dashed lines. Gray dashed circles, nucleoli. Scale bar = 10 μm. **(B)** Quantification of cells with fully segregated nucleoli (nucleolar caps) as in (A). Red lines = mean, ±SD is shown (N=3, >400 cells). **** p<0.001, paired two-sided Student’s t-test compared to DMSO control. **(C)** Schematic of the rDNA damage dropout screen. hTERT-RPE1 *p53*^-/-^ cells were transduced with the TKOv3 library and selected for using puromycin. Cells were divided into 2 technical replicates and treated with DMSO or Shield-1+4-OHT over the course of 3 weeks. Timepoint 16 (T16) and 24 (T24) were used for sgRNA sequencing and screen analysis. Illustration of genes whose knockout results in cell death is shown on the far right. **(D)** NormZ analysis of CRISPR-Cas9 dropout screen, cells at T16 exposed to I-PpoI induced rDNA damage as in (C). Purple, genes of protein UFMylation pathways with NormZ score. Black, genes considered non-significant hits. Orange, N=934 genes sensitizing cells to rDNA DSBs (NormZ ≤1.5). **(E)** Protein UFMylation pathway. Cleavage of pro-UFM1 by UFSP1/2 produces mature UFM1 with exposed C-terminal glycine residue. UFM1 is conjugated to UBA5 via a thioester bond, transferred to UFC1 via a transthioation reaction, and then modified on a substrate lysine residue in a process which requires the E3-like protein UFL1. UFL1-binding partners (DDRGK1 and CDK5RAP3) catalyse UFMylation and provide substrate specificity. UFMylation is reversible via UFSP1/2 cleavage. **(F)** NormZ analysis of CRISPR-Cas9 dropout screen, cells at T16 exposed to I-PpoI induced rDNA damage as in (C). Green, genes of chromatin remodellers involved in H3.3 deposition with NormZ score. Black, genes considered non-significant hits. Orange, N=934 genes sensitizing cells to rDNA DSBs (NormZ ≤1.5). **(G)** NormZ analysis of CRISPR-Cas9 dropout screen, cells at T16 exposed to I-PpoI induced rDNA damage as in (C). Genes known to regulate rDNA DSB repair or rDNA transcriptional silencing in grey alongside their NormZ score. Orange, N=934 genes sensitizing cells to rDNA DSBs (NormZ ≤1.5). Blue, N= 3 genes providing resistance to rDNA DSBs (NormZ ≥6).

We then used this cell line to conduct a genome-wide CRISPR-Cas9 loss-of-function genetic screen using the Toronto Knockout Version 3 (TKOv3) sgRNA library to identify genes essential for rDNA DSB repair (Figure 1C)^43,44^, anticipating sgRNAs targeting genes essential for rDNA DSB repair to be underrepresented in I-PpoI-induced cells compared to uninduced cells as they would be unable to repair rDNA lesions leading to apoptosis. To that end, we transduced cells with the TKOv3 sgRNA library and subsequently split cells into two groups for acute treatment with either DMSO or Shield-1+4-OHT at the pre-determined LD_20_ concentration (Figure S1E). Single guide RNAs present in the different cell populations were amplified, sequenced, and analyzed using the DrugZ algorithm to identify chemical genetic interactions and the BAGELv2 gene essentiality algorithm^45,46^. We focussed our analysis on the T16 timepoint based on performed quality control measurements, in which we generated precision-recall curves and assessed read counts of the core and non-essential genes with a minimum coverage of 200X (Figure S2A-C).

To determine genes whose functions are essential for rDNA DSB repair, we applied a NormZ cut-off of ≤ -1.5 to identify gene knockouts that sensitized cells to rDNA DSBs, resulting in an identification of 934 genes with a p-value ≥ 0.05 (Figure 1D, E; gray dashed line). Gene ontology (GO) enrichment analyses revealed a significant enrichment in genes associated with ubiquitin-like protein transferase activity and protein modification processes (Figure S2C-E). A detailed analysis of our data identified components of the protein UFMylation pathway as being essential for rDNA DSB repair (Figure 1D, Figure S3A).

Similar to ubiquitin, the Ubiquitin Fold Modifier-1 (UFM1), a ubiquitin-like modifier, is covalently transferred posttranslationally to the primary amine of a lysine residue in target proteins via the coordinated action of three enzymes: a UFM1-activating E1, a UFM1-conjugating E2, and a UFM1-ligase E3, together with several components of the UFMylation machinery (Figure 1E)^29,33,47^. UFMylation is reversible via cleavage by UFSP1/2, two enzymes also responsible for biogenesis of UFM1^29,48,49^. Several proteins that may provide UFMylation substrate specificity such as DDRGK1 and CDK5RAP3 have also been identified as UFL1 binding partners^50–52^. However, unlike ubiquitination, only one protein for each step of the cascade has so far been described^29,33^.

While protein UFMylation has been linked to a number of cellular stress response pathways as well as the control of interconnected cellular processes such as ER homeostasis, viral infection and immune signalling, ribosome stalling and quality control, and telomer maintenance, and recent work has also suggested a potential role for UFMylation in nuclear DNA damage response, so far UFMylation has not been implicated in ribosomal DNA repair^29,33^. Our screen identified all genes encoding for the components of UFMylation machinery as essential sensitizers for rDNA damage, including for the E1 enzyme UBA5, the E2 enzyme UFC1, the E3-like enzyme UFL1, the maturation enzyme UFSP2 and its interactor C1orf27 (ODR4), as well as those encoding for UFL1-binding adapter proteins DDRGK1 and CDK5RAP3, and UFM1 (Figure 1D, F), suggesting that the UFMylation pathway is critical for functional ribosomal DNA damage response and repair.

We furthermore identified several genes encoding for key chromatin organizers as essential for rDNA DSBs repair, in particularly, those important for histone H3/H3.3 disposition on chromatin, including *HIRA, CABIN1* and *UBN1* as well as *TLK-1* and *-2, ATRX* and *DAXX* (Figure 1F)^53–61^. Of these, only ATRX has previously been linked to rDNA heterochromatin formation and stability^54^.

In addition, members of the HUSH complex (*FAM208A, MORC2*, and *MPHOSPH8)* and the effector H3K9Me3 methyltransferase *SETDB1* were identified as essential for rDNA DSBs repair in our screen (Figure 1G, Figure S3B), in line with previous studies^11,30^, as well as a number of known rDNA chromatin regulators such as *RPA2, NBN*, and *JMJD6*, the latter two expressing known TCOF1 interactors, were also found to be essential in our screen, validating its robustness^40,54,62–64^. Notably, we did not identify *TCOF1* in our screen as it is a known essential gene in hTERT-RPE1 cells and thus showed already significantly reduced sgRNA counts at our initial collection timepoint, even prior to induction of rDNA DSBs (Table S1)^65^; these same was observed for *TOPBP1* which is also known to be essential in hTERT RPE1 cells^66^. Moreover, although sensitizers, we did not identify genes of known downstream DNA damage response factors (*e.g*., *BRCA1, 53BP1, BRCA2, RAD51*) as essential for rDNA DSB repair, suggesting either that repair might occur via multiple parallel pathways and/or that genes identified here, encode for factors that comprise essential earlier steps in the rDNA damage response, upstream of these processes.

Using a NormZ cutoff of ≥ 6, we also identified three genes associated with resistance to rDNA DSBs, namely *FKBPL, ZCCHC14, NXT1* (Figure 1D,F,G, blue; Table S2). *NXT1* showed the highest resistance score, possibly due to its function in mRNA export^67^, as the loss of I-PpoI mRNA export would likely preclude its protein production and stabilization in the cytoplasm, and, consequently, its nuclear import and targeting to rDNA for DSB formation, resulting in resistance to cell death.

Taken together, our loss-of-function genetic screen identified, for the first time, the protein UFMylation machinery as critical for rDNA DSB repair in human cells, in addition to chromatin regulation factors. Moreover, our data suggests that while parallel and redundant pathways exist for the repair of ribosomal DNA DSBs, processes initiating rDNA repair (*i.e*., rDNA segregation and nucleolar cap formation, transcriptional silencing) require distinct signaling pathways and factors, including UFMylation and chromatin remodelers.

### The protein UFMylation pathway and HUSH complex are important for nucleolar segregation and cell survival upon rDNA damage

To further validate the results of our screen and determine the importance of UFMylation on cell survival upon rDNA damage, we assessed clonogenic survival in hTERT-RPE1 *p53-/-* Cas9 DD-HA-ER-I-PpoI cells after knock-out of the major components of the UFMylation pathway (Figure 2A). Cells transduced with sgRNAs targeting *UFM1, UFSP2, UBA5, UFL1* or *DDRGK1* had higher sensitivity to I-PpoI-induced rDNA damage when compared to control cells (∼ 50%, 53%, 62%, 57% and 70% decrease in cell viability for *UFM1, UFSP2, UBA5, UFL1* and *DDRGK1* compared to sg*LacZ* control cells, respectively), confirming the importance of the UFMylation pathway for cell viability in response to rDNA DSBs (Figure 2A, B).

**Figure 2.**
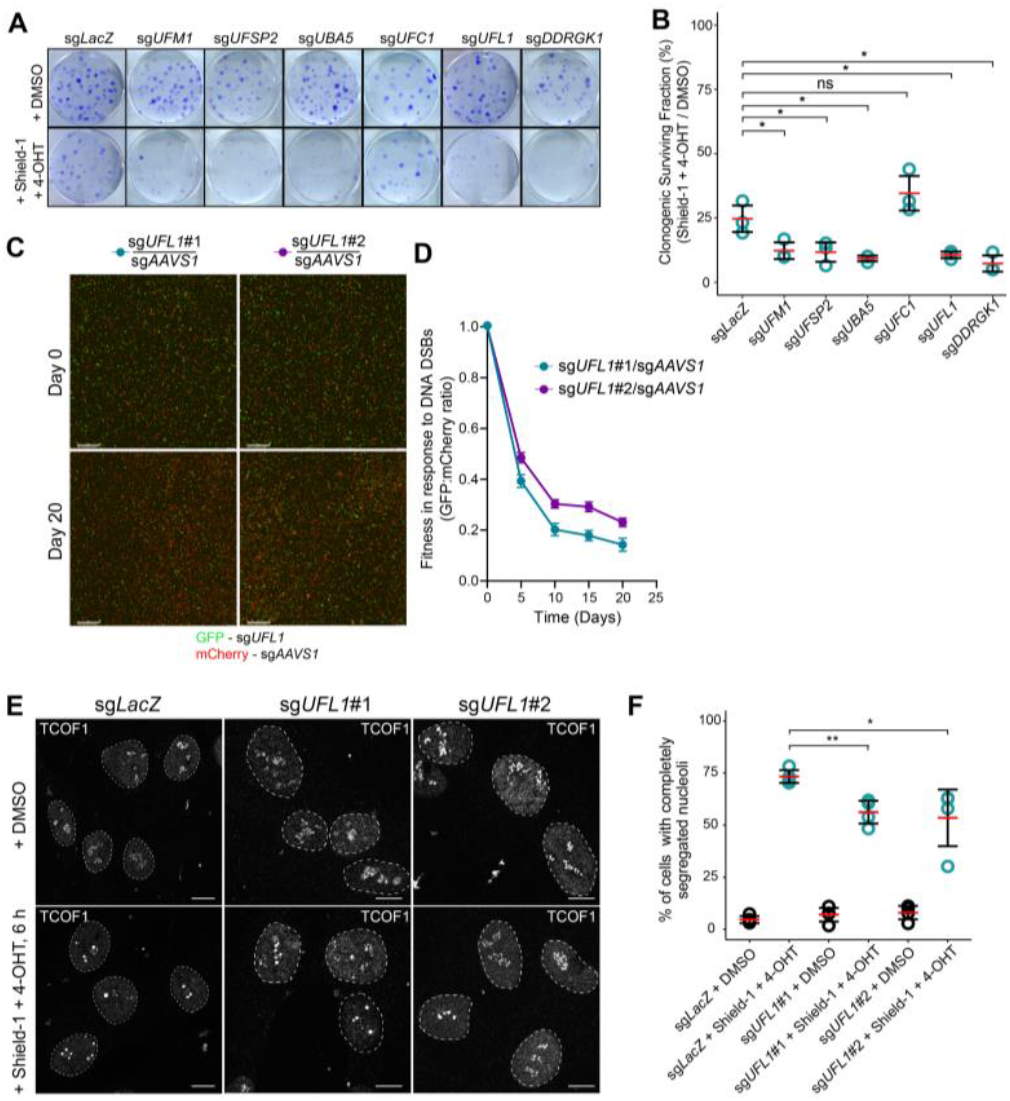
The UFMylation pathway is required for cell viability and survival in response to rDNA DSBs. **(A)** Clonogenic survival assay of hTERT-RPE1 *p53*^-/-^ DD-HA-ER-I-PpoI cells transduced with sgRNAs targeting different genes of the UFMylation pathway. Cells treated with DMSO or Shield-1+4-OHT every 3 days for a total of 10-14 days then stained with crystal violet. **(B)** Quantification of the clonogenic surviving assay in (A). Y-axis: Shield-1+4-OHT/DMSO normalized clonogenic surviving fraction. Red lines = mean, ±SD is shown (N=3); ns, no significance, * p<0.05, unpaired two-sided Student’s t-test compared to sg*LacZ*. **(C)** Competitive growth assays of hTERT-RPE1 *p53*^-/-^cells expressing DD-HA-ER-I-PpoI transduced with two independent *UFL1* targeting sgRNAs (co-expressing GFP) compared to a control *AAVS1* targeting sgRNA (co-expressing mCherry). Cells plated at 1:1 ratio and passaged every 3 days. Cells treated with Shield-1+4-OHT 24 hr after plating. Cells imaged every 24 hr over 20 days. N=3. Scale bar = 800 μm. **(D)** Quantification of fitness (GFP:mCherry ratio) of cells shown in (C). N=3, error bars represent SD. **(E)** Immunofluorescent analysis of nucleolar cap formation in hTERT-RPE1 *p53*^-/-^ Cas9 DD-HA-ER-I-PpoI cells transduced with *UFL1*-targeting gRNAs. Cells were treated with DMSO or Shield-1+4-OHT for 6 hr to induce nucleolar segregation. Scale bar = 10 μm. Dashed lines outline nucleus. **(F)** Quantification of nucleolar caps as shown in (E). Red lines = mean, ±SD is shown (N=4, >250 cells). * p<0.05, ** p<0.01, unpaired two-sided Student’s t-test compared to sg*LacZ*.

Since the E3-like protein UFL1 has been suggested to interact with the MRN complex, localize to DNA damage foci, and regulate MRE11 protein interactions on nuclear DNA, we further investigated its importance for rDNA DSB repair^34,35,37^. To assess if UFL1 is required for cell fitness in response to rDNA DSBs, we carried out a two-colour competitive growth assay using a control sgRNA targeting the *AAVS1* locus co-expressed with mCherry or two independent sgRNAs targeting *UFL1* co-expressed with GFP in cells treated with Shield-1+4-OHT every three days for continued rDNA damage (Figure 2C). Using quantification of the ratio between GFP:mCherry as readout of fitness, we observed that after 20 days only ∼20-30% of cells transduced with *UFL1-* targeting sgRNAs remained in the cell population, suggesting that loss of UFL1 negatively affected cell fitness under rDNA damage conditions compared to control cells (Figure 2D).

As we did not observe general DNA repair proteins as being essential for rDNA repair in our screen, we hypothesized that the identified genes might encode factors that affect early steps of rDNA repair, including nucleolar segregation and transcriptional silencing. We therefore assessed whether UFL1 is required for nucleolar segregation in response to rDNA DSBs. To that end, we analyzed levels of nuclear cap formation in cells transduced with either *UFL1* sgRNAs or control *LacZ* sgRNA 6 hours after induction of rDNA damage using fluorescent microscopy and TCOF1 as a marker for nucleolar segregation (Figure 2E). We observed that full formation of nucleolar caps and nucleolar segregation was reduced to ∼55% in cells lacking UFL1, compared to ∼73% in control cells (Figure 2F). As UFL1 is the only known UFM1 E3 ligase, our data suggests that loss, or even reduction, of UFMylation in the absence of UFL1 affects efficiency of nucleolar segregation and cap formation, furthermore implicating UFMylation as an important upstream regulator of rDNA repair.

As previous data suggested a role for the HUSH complex in transcriptional silencing of rDNA^11^, we also tested the importance of *FAM208A* and *SETDB1* for cell viability under rDNA damage conditions. Once again, we conducted clonogenic survival assays in hTERT-RPE1 *p53-/-* Cas9 DD-HA-ER-I-PpoI cells after knock-out of *FAM208A* and *SETDB1* via with sgRNAs under DD-HA-ER-I-PpoI induced rDNA damage (Figure S3C). Depletion of both *FAM208A* and *SETDB1* resulted in a significant decrease in colonies compared to control cells (∼72-96% and ∼43-52% decrease in cell viability, respectively) (Figure S3D), suggesting that, while both are involved in rDNA DSB repair, the HUSH complex may play a significant role in the early steps of the pathway.

Taken together, our results confirm a critical importance of the UFMylation machinery and HUSH complex for cell viability in response to ribosomal DNA damage. Moreover, while we cannot rule out that UFM1 has more than one E3 ligase, which has yet to be identified, the significant decrease in nucleolar cap formation upon loss of UFL1 under damage conditions suggests that UFL1 and UFMylation are essential for efficient nucleolar segregation and rDNA DSBs repair.

### Transcriptional rDNA silencing and ATM activation is reduced in *UFL1*-deficient cells

Transcriptional silencing of rDNA is a pre-requisite for rDNA re-localization into nucleolar caps^11,14^. The overall decrease in nucleolar cap formation upon UFL1 knock-down suggest that UFL1/UFMylation might affect nucleolar segregation directly or as a consequence of a reduction in transcriptional silencing of rDNA. To further investigate the downstream consequences of loss of UFL1 and effects of UFMylation on rDNA damage response, we generated a *UFL1* knockout (KO) in hTERT-RPE1 WT cells (Figure S4A). To test if loss of UFL1/UFMylation affected rDNA silencing, we analyzed nucleolar EU-staining as readout of active rRNA transcription using fluorescent microscopy and TCOF1 as marker for nucleolar cap formation (Figure 3A)^68,69^. Quantification of EU showed that overall *UFL1 KO* cells had already lower basal levels of rRNA transcription (∼22%) compared to DMSO control cells prior to damage (Figure 3B), suggesting that UFL1/UFMylation affects rDNA transcription. Induction of rDNA DSBs by I-PpoI resulted in ∼30% global reduction of rRNA transcription in *LacZ* transduced cells, in line with previous observations^10–13^, while in *UFL1* KO cells only a partial silencing of rRNA transcription (∼15%) was observed upon damage (Figure 3A, arrowheads; Figure 3B). In line with EU staining, northern analysis of 47S pre-rRNA levels confirmed both the reduction in basal levels of 47S and rDNA transcription in undamaged *UFL1* KO cells (Figure 3C, lanes 1 and 5; Figure 3D; ∼28%) and partial silencing of rDNA transcription upon induction of DSBs in *UFL1* KO compared to controls (Figure 3C; Figure 3D; ∼43% and ∼56%, respectively).

**Figure 3.**
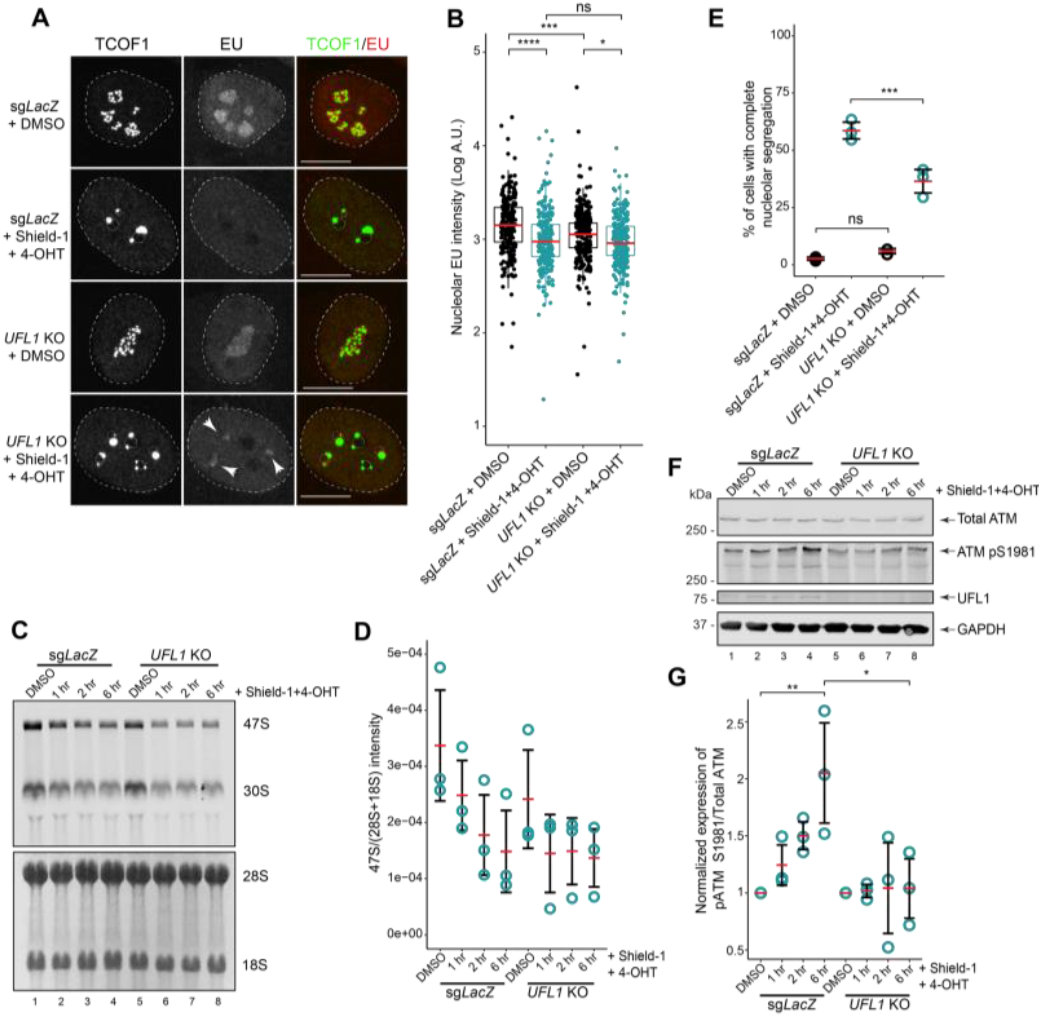
*UFL1*-deficient cells have reduced rRNA transcription, nucleolar segregation, and ATM activation. (**A**) EU-staining in wild-type hTERT-RPE1 *p53*^-/-^ Cas9 DD-HA-ER-I-PpoI or *UFL1* KO cells + DMSO or Shield-1+4-OHT for 6 hr to induce nucleolar segregation. Maximum projection images of single cells are shown. DAPI, nuclei outlined with white dashed lines; grey dashed circles, nucleoli. Scale bar = 10 μm. (**B**) Nuclear EU signal quantification as shown in (A). ns, no significance, ** p<0.01, *** p<0.001, **** p<0.0001, paired and unpaired two-sided Student’s t-test. Red bars = median. N=3, >230 cells. (**C**) Northern blot of total mature (28S/18S) and precursor (47S/30S) rRNA in wild-type hTERT-RPE1 *p53*^-/-^ Cas9 DD-HA-ER-I-PpoI or *UFL1* KO cells + DMSO or Shield-1+4-OHT. Staining with methylene blue and a probe for the 5’-ETS. (**D**) Northern quantification, 47S rRNA intensity over time as normalized to total combined 28S and 18S rRNA intensity as in (C). Red bars = mean. N=3. (**E**) Quantification of cells with full nucleolar segregation as in (A). ns, no significance, *** p<0.001, one-way ANOVA with Tukey’s post-hoc test. N=3, >230 cells. (**F**) Western analysis, total ATM, pATM S1981, UFL1 and GAPDH protein levels in wild-type hTERT-RPE1 *p53*^-/-^ Cas9 DD-HA-ER-I-PpoI or *UFL1* KO cells + DMSO or Shield-1+4-OHT. (**G**) Quantification of pATM-S1981 over total ATM levels as in (F). * p<0.05, N=3.

Taken together, our data suggest that UFL1/UFMylation affects, and potentially regulates, rDNA transcription, including its silencing upon rDNA damage. The reduced efficiency of rDNA silencing in *UFL1* KO cells is likely a consequence of the overall reduction of active rDNA genes and reduced rRNA transcription.

Since transcriptional silencing of rDNA is required for nucleolar segregation^11,12^, we also determined the proportion of *UFL1* KO cells with complete nuclear segregation. After rDNA DSBs induction ∼37% of *UFL1* KO cells exhibited solely nucleoli that went complete nucleolar segregation compared to ∼60% in control cells (Figure 3E), while the majority of cells displayed a mixture of full, but mostly partially, and non-segregated nucleoli, confirming our previous results that loss of UFL1/UFMylation prevents efficient nucleolar segregation (Figure 2E); the observed reduction in nucleolar segregation was not linked to changes in the cell cycle as no discernable differences in cell cycle kinetics between *LacZ* and *UFL1* KO cells in DMSO or upon induction of rDNA DSBs were observed (Figure S4B).

rDNA DSB-induced repression of RNA Pol I transcription is dependent on the major DNA repair serine/threonine kinase ATM ^5,10,12,13,24,70^. Previously, UFL1 was suggested to be important for ATM activation following DSB induction on chromatin via ionizing irradiation (IR)^34^. To test whether the observed reduction of rDNA transcriptional silencing in *UFL1* KO cells upon damage was due to altered ATM activation, we analyzed total and Ser1981-phosphorylated ATM at 1, 2, and 6 hours after rDNA DSB induction (Figure 3F). While *LacZ* control cells showed ∼2-fold increase in Ser1981-phosphorylated ATM after 6 hours, no substantial change in ATM phosphorylation was observed in *UFL1* KO cells, suggesting that the low levels of rDNA transcriptional silencing may be a consequence of insufficient ATM activation and that UFL1/UFMylation is likely important for ATM activation, not only during DSB in the nucleus but also along the ribosomal DNA damage response pathway (Figure 3F, G).

Previous work further determined that recruitment of BRCA1 and 53BP1, two key homologous-recombination and homologous-end-joining repair proteins, respectively, to rDNA/nucleoli in cells lacking TCOF1, where nucleolar cap formation is impaired, was significantly reduced^12,14^. A similar reduction in the recruitment of both proteins to nuclear DSB sites had recently been observed in UFL1-depleted cells^34^. Since loss of UFL1 caused a significant reduction in nuclear cap formation (Figure 2E, F) as well as rDNA transcriptional silencing (Figure 3A-D), we tested if UFL1 was important for the recruitment of these DNA repair proteins to nucleoli/rDNA by assessing co-localization of 53BP1 and BRCA1 with TCOF1 in undamaged and I-PpoI damaged cells^71–73^. While 53BP1 and BRCA1 was found co-localized with TCOF1 in nucleoli with fully segregated nucleolar caps, which were present in fewer cells, in line with our previous observations (Figure 2E, F), reduced or no co-localization of BRCA1 and 53BP1 with TCOF1 was observed in non-or partially segregated nucleoli in *UFL1* KO cells upon damage induction (Figure S4C, D; arrowheads), suggesting that UFL1/UFMylation is not required for the downstream recruitment of DNA repair proteins to segregated nucleolar caps for rDNA repair (Figure S4C, D; blue arrowheads) but rather during steps preceding nucleolar cap formation and rDNA segregation.

Taken together, our results suggest that UFL1/UFMylation plays a critical role in the early steps of the rDNA damage response, for efficient activation of ATM, rDNA transcriptional silencing, and nucleolar segregation; whether the observed effects on each of these steps is a consequence of ineffective ATM activation, or whether they are each modulated by UFL1/UFMylation individually, remains to be determined.

### UFL1 and DDRGK1 associate with TCOF1 in nucleolar caps upon rDNA damage

UFL1 was shown to localize to sites of IR-induced DSBs on chromatin^34^, and our data observed a critical importance of UFL1/ UFMylation for early steps of rDNA damage response. To further determine whether UFL1 was localized to nucleolar caps upon induction of rDNA DSBs in hTERT-RPE1 cells, we determined its association with TCOF1 as a known critical regulator of nucleolar cap formation and early rDNA damage response ^12,14,16,74^.

While TCOF1 was shown to function primarily in rDNA transcription and early ribosome biogenesis^75,76^, upon rDNA damage, phosphorylation of TCOF1 by ATM facilitates recruitment of the MRE11-RAD50-NBS1 (MRN) complex to nucleoli^12,14,16,74^. These steps were shown to be instrumental for ATM/ATR activation, RNA Pol I repression, subsequent nucleolar cap formation and rDNA repair^12,14–16^. Cells lacking TCOF1 are unable to form nucleolar caps or efficiently recruit downstream DNA repair proteins (*e.g*., BRCA1/2, 53BP1) to the nucleolus^12^. Its pivotal function in nuclear cap formation and early steps of rDNA repair made TCOF1 an ideal bait protein for the isolation and analysis of an early rDNA repair proteome and to assess the presence of UFL1 and/or other proteins of the UFMylation machinery in nucleolar caps.

We therefore stably integrated TCOF1-eGFP in the same hTERT-RPE1 cells utilized in our CRISPR-Cas9 genetic screen. Immuno-fluorescent microscopy showed punctate nucleolar localization of TCOF1-eGFP in control cells, while addition of Shield-1+4-OHT for rDNA DSBs induction resulted in re-localization of TCOF1-eGFP into nucleolar caps after 6 hours, where it co-localized with an early key DNA repair factor, NBS1, a component of the MRN complex, as expected (Figure 4A; Figure S3E)^16,74^. Moreover, nucleolar segregation was observed in ∼80% of cells 6 hr after treatment with Shield-1+4-OHT, compared to less than 10% in DMSO-treated cells (Figure 4B).

**Figure 4.**
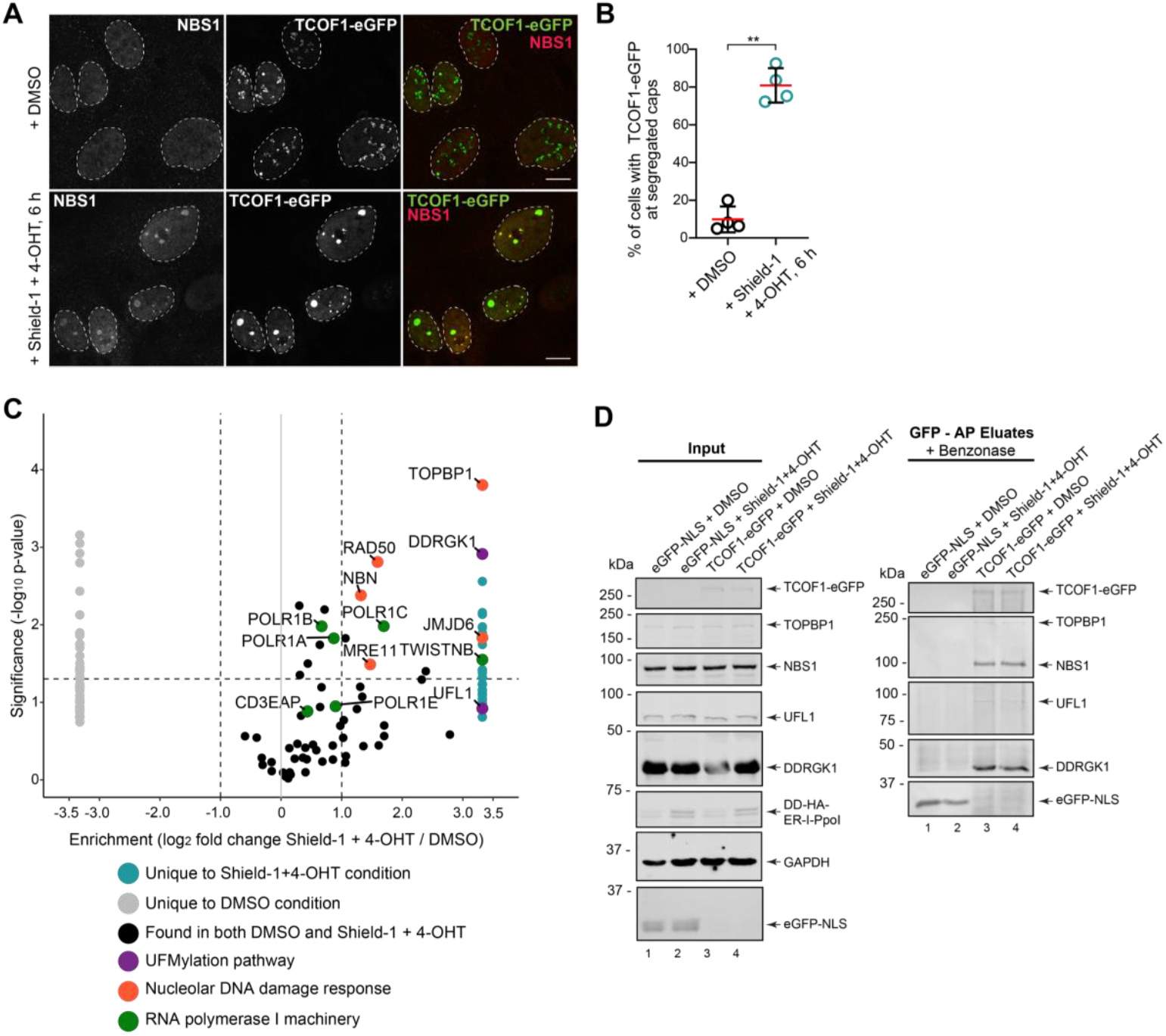
UFL1 and DDRGK1 form a complex with TCOF1 at damaged rDNA. **(A)** Microscopy of hTERT-RPE1 *p53*^-/-^ Cas9 cells stably expressing DD-HA-ER-I-PpoI and TCOF1-eGFP. Cells + DMSO or Shield-1+4-OHT for 6 hr. Scale bar = 10 μm. **(B)** Quantification of TCOF1-eGFP expressing cells with segregated nucleoli (nucleolar caps) as in (A). Red lines = mean, ±SD are shown as determined by affinity-purification mass spectrometry in cells + DMSO or Shield-1+4-OHT for 6 hr. TCOF1-eGFP or eGFP-NLS purified from whole cell extracts using GFP-nanobody conjugated Dynabeads and analyzed by MS. Proteins found exclusively in either the DMSO control or Shield-1+4-OHT conditions were assigned an arbitrary log2-fold change of -3.5/3.5, respectively and labelled in grey or teal. Horizontal lines -log_10_p =1.301 (p=0.05). Vertical lines log_2_ fold-change of cells + Shield-1+4-OHT/DMSO of 1. N=3 biological replicates. **(D)** Affinity purification western blot of TCOF1-eGFP associated proteins as in (C). Cells expressing eGFP-NLS or TCOF1-eGFP + DMSO or Shield-1+4-OHT for 6 hr. Whole cell extract inputs on the left; eluates of TCOF1-GFP complexes from benzonase-treated lysates shown on the right.

Affinity-purification (AP) and analysis of TCOF1-eGFP-associated complexes by semi-quantitative mass spectrometry (MS) in undamaged conditions identified several ribosome biogenesis factors in accordance with its function in rRNA transcription^68^ (Table S8). Analysis of complexes purified 6 hours after I-PpoI induction revealed an enrichment of several known DNA repair proteins, including the MRE11, RAD50, NBS1 (MRN complex), TOPBP1, and JMJD6, all known TCOF1 interactors (Figure 4C, log2 fold change ≥ 1 and p-value ≥ 0.05 relative to undamaged cells; Table S8)^12,16,40,74^. Moreover, in line with a role for UFL1/UFMylation in rDNA repair, we identified the UFM1-E3-ligase UFL1 and its direct interacting partner, DDRGK1 (UFBP1), which structurally complements UFL1 and promotes both its stability and E3 ligase activity, in cells after damage but not undamaged conditions (Figure 4C; Table S8)^33^; neither proteins had so far been reported to localize to nucleolar caps upon rDNA damage. We further identified PHF6 under these conditions, a transcriptional regulator that was previously shown to associate with ribosomal RNA promoters and to suppress ribosomal RNA (rRNA) transcription^77^, and several subunits of RNA polymerase I (POLR1A/B/C/E, CD3EAP, TWISTNB), suggesting that RNA Pol I may remain associated with rDNA during repair in nucleolar caps, while rDNA transcription is repressed (Figure 4C; Table S8).

Components of the MRN complex were also purified with TCOF1 in undamaged conditions, which is likely due to continued basal damage in those cells as we observe minimal formation of nucleolar caps (∼ 4%) in cells in non-damage conditions (Figure 4B). Association of UFL1 and DDRGK1 with TCOF1 in nucleolar caps upon rDNA DSB induction was further confirmed by western blotting in eluates after benzonase treatment, indicating moreover that their association with TCOF1 complexes is chromatin-independent (Figure 4D). This suggest that UFL1 and DDRGK1 are likely targeted to nucleolar caps via protein interactions and/or might interact with and UFMylate TCOF1 itself or its interaction partners.

Presence of UFL1 and DDRGK1 in nucleolar caps upon damage was also observed by fluorescent microscopy (Figure 5). Displaying a predominately diffused nuclear localization in undamaged cells (Figure 5A, Figure S5A, B), upon I-PpoI induction, UFL1 formed discrete nuclear foci most of which colocalized with TCOF1 in nucleolar caps (Figure 5A-C; Figure S5A, B). As DDRGK1 was significantly enriched in TCOF1-associated complexes upon rDNA damage induction (Figure 4C) and has been reported to form a heterodimer with UFL1^32^, we also determined DDRGK1 localization. Similar to UFL1, DDRGK1 was displayed a mostly diffuse nuclear localization in undamaged cells (Figure 5D, Figure S5C, D); induction of I-PpoI resulted in the increased formation of DDRGK1 foci many of with co-localized with TCOF1 to nucleolar caps (Figure 5D-F; Figure S5C, D). Some foci formation for both proteins was also observed outside nucleoli, supporting previously reported recruitment of UFL1 to DSB sites on chromatin, and in line with known I-PpoI cleavage sites elsewhere in the genome ^10,13,42,78^.

**Figure 5.**
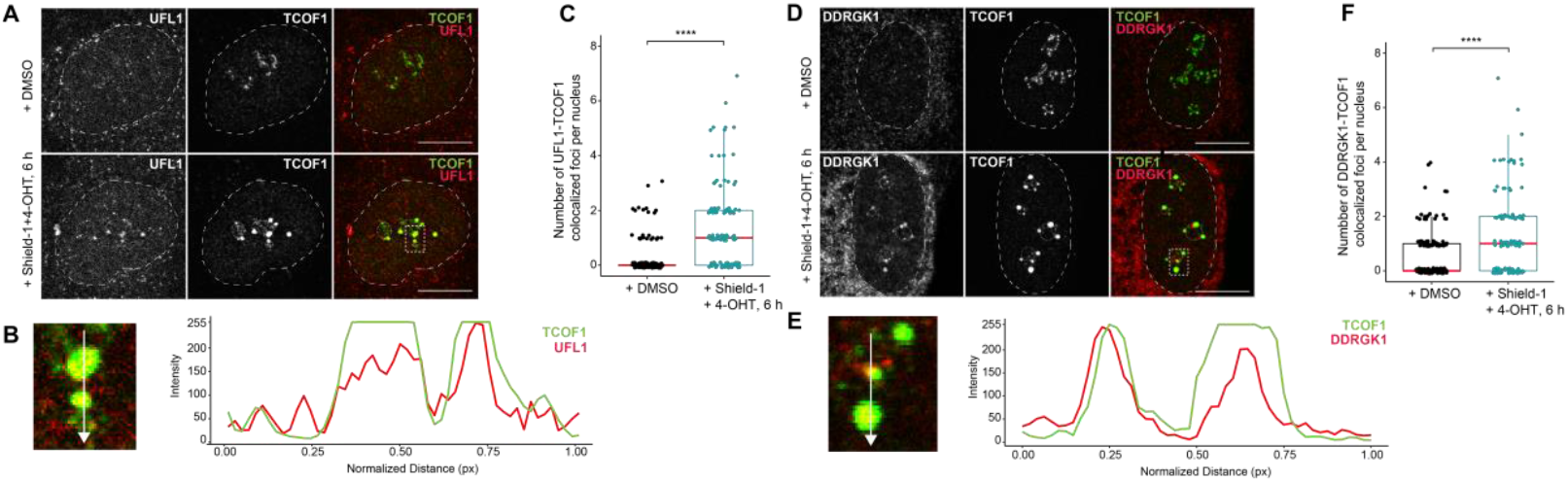
UFL1 and DDRGK1 colocalize with TCOF1 in segregated nucleolar caps. **(A)** Microscopy analysis of hTERT-RPE1 *p53*^-/-^ Cas9 DD-HA-ER-I-PpoI cells + DMSO or Shield-1+4-OHT for 6 hr. Nuclei and nucleoli outlined with dashed lines. Scale bar = 10 μm. (N=3, >125 cells). Single Z-stack slices are shown. **(B)** Zoom of nucleolar cap from (D). White arrow represents the line scan path of relative signal intensity for UFL1 (red) and TCOF1 (green). **(C)** Quantification of co-localized UFL1-TCOF1 foci from (A). Red lines = median. **** p<0.0001, paired two-sided Student’s t-test compared to DMSO control. (N=3, >125 cells) **(D)** Microscopy analysis of hTERT-RPE1 *p53*^-/-^ Cas9 c]DD-HA-ER-I-PpoI cells + DMSO or Shield-1+4-OHT for 6 hr. Scale bar = 10 μm. Single Z-stack slices are shown. Nucleus and nucleoli indicated by dashed lines. **(E)** Zoom of nucleolar cap from (D). White arrow represents the line scan path of relative signal intensity for DDRGK1 (red) and TCOF1 (green). **(F**) Quantification of co-localized DDRGK1-TCOF1 foci from (C). Red lines = median. *** p<0.001, **** p<0.0001, paired two-sided Student’s t-test compared to DMSO control. (N=3, >125 cells).

UFMylation has been implicated in several cytoplasmic stress response pathways^33^. To determine whether UFL1 and DDRGK1 translocate from the cytoplasm into the nucleus upon rDNA DSB induction, we conducted subcellular fractionation and western blotting in wild-type, *UFL1* KO and *DDRGK1* KO cells (Figure S6). *UFL1* KO cells showed overall reduced protein levels of DDRGK1, while *DDRGK1* KO cells had overall reduced levels of UFL1 in both the nuclear and cytoplasmic fractions compared to wildtype (Figure S6A), consistent with recent findings suggesting that UFL1 and DDRGK1 heterodimerize to stabilize and activate UFL1 function^32^. Moreover, no increase in nuclear UFL1 or DDRGK1 protein levels was observed upon rDNA DSB induction (Figure S6A, lanes 4, 8, and 12), suggesting that the increase in nucleolar UFL1 and DDRGK1 foci after I-PpoI induction is likely due to re-localization of an already existing nuclear and/or nucleolar pool of UFL1 and DDRGK1. This is in line with nuclear staining and low number of nuclear and nucleolar foci of UFL1 and DDRGK1 under non-damage conditions (Figure 5A, B; Figure S5B, D) and previous suggestion of DDRGK1 as potential nucleolar protein^79^.

Taken together, our data suggest that, in response to rDNA damage, UFL1 and DDRGK1 localize to nucleolar caps where they colocalize and interact with TCOF1-associate complexes in a chromatin-independent manner, further suggesting a role for not only UFL1 but also DDRGK1 and UFMylation in the early rDNA damage response pathway. Furthermore, our data also indicates that while UFL1 and DDRGK1 heterodimerize, they may exist independently of one another in the nucleus and nucleolus; the trigger and regulation of UFL1/DDRGK1 dimerization remains so far unknown.

### Chromatin segregation and DNA repair factors are UFMylated in response to DNA damage

UFMylation has previously been identified as regulator of cytoplasmic stress response pathways, particularly ER proteostasis^33^. An additional role in gene expression and maintenance of genome integrity has also been suggested as UFL1 was shown to colocalize with DSB on chromatin and UFMylate MRE11 to increase ATM activation along genomic DNA damage response pathways^37^. Moreover, DDRGK1, had been suggested to localize to the nucleolus^79^, which was confirmed by our observations, with a significant increase in nucleolar DDRGK1 foci in response to rDNA damage, upon which it was observed to colocalize with TCOF1 in nucleolar caps (Figure 4C; Figure 5C, D). While our data suggest that protein UFMylation affects early steps of the rDNA damage response pathway, its nuclear and nucleolar targets upon rDNA DSB induction, besides MRE11, remain unknown.

To determine nuclear/ nucleolar UFMylated targets in undamaged conditions and in response to rDNA DSBs, we generated an hTERT-RPE1 cell line stably expressing His_10_-UFM1ΔSC. The C-terminal deletion of Ser-Cys (termed ΔSC) has previously been used to bypass the need for UFSP1/2-mediated biogenesis of pro-UFM1 into a conjugatable form of UFM1, exposing the reactive C-terminal glycine residue and allowing for enrichment of UFMylated proteins within cells^49,80,81^. We purified His_10_-tagged UFM1ΔSC-conjugated proteins from purified nuclei lysed under denaturing, non-reducing conditions (Figure 6A) and determined nuclear UFMylated proteins using semi-quantitative mass spectrometry (Figure 6C). Confirmation of efficient UFMylation and isolation of UFM1-conjugated proteins from nuclei was carried out by western (Figure 6B), and no changes in UFMylation levels of highly abundant proteins, including RPL26 (uL24), which may also be UFMylated in the nucleolus^29–31,80,82^, were detected between non-damage and damage conditions, while UFM1 and its associated proteins were efficiently isolated.

**Figure 6.**
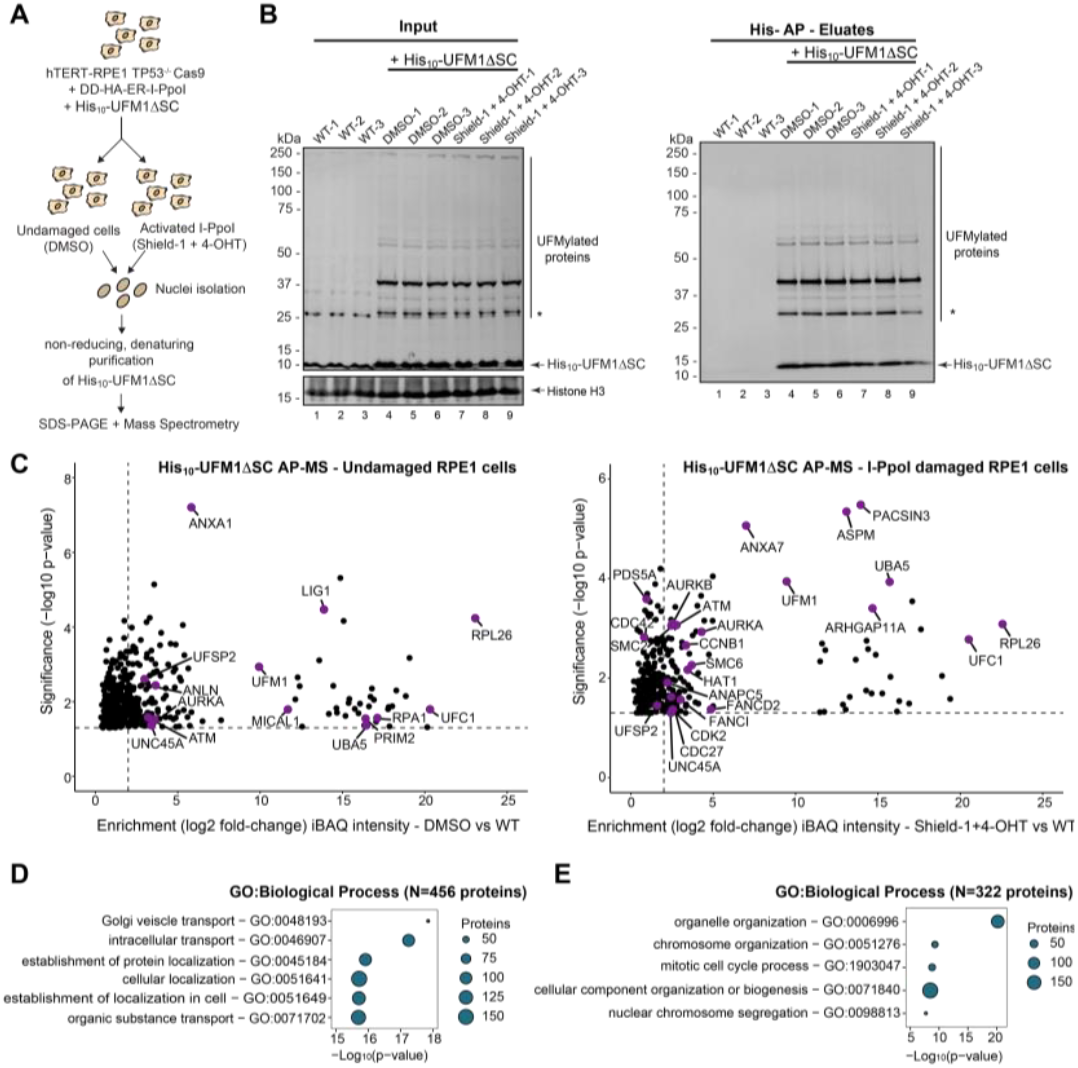
Identification of UFMylated proteins in response to rDNA DSBs. (**A**) Nuclei isolation and purification of His_10_-tagged UFMylated proteins. hTERT-RPE1 p53-/-Cas9 DD-HA-ER-I-PpoI cells stably expressing His_10_-UFM1ΔSC + DMSO or Shield-1+4-OHT for 6 hr prior to nuclei isolated in a hypotonic buffer and with 8 M urea. His_10_-UFM1-conjugated proteins were isolated via magnetic Ni-NTA resin and analyzed by MS and western. (**B**) Western analysis of hTERT-RPE1 p53-/-Cas9 DD-HA-ER-I-PpoI His_10_-UFM1ΔSC nuclear extracts. Inputs on the left, eluates of His_10_-UFM1-conjugated proteins on the right. N=3 biological replicates. (**C**) Scatterplot of nuclear/nucleolar UFMylated proteins identified using MS as in (A). Proteins significantly enriched over wild-type cells (p≤0.05) are shown. Horizontal lines -log10p=1.301 (p=0.05). Vertical lines log2 fold-change iBAQ intensities = 2. Proteins identified + DMSO on the left, + Shield-1+4-OHT on the right. Key proteins are in purple. (**D**) The top six GO-term biological process enrichment of all identified nuclear/nucleolar UFMylated proteins (p≤0.05) purified in + DMSO conditions as in (C). N=456 proteins in total. (**E**) The top 5 GO-term biological process enrichment of all identified nuclear/nucleolar UFMylated proteins (p≤0.05) purified in + Shield-1+4-OHT conditions as in (C). N=322 proteins in total.

UBA5 and UFC1, the E1 and E2 catalyzing and conjugation enzymes for UFM1 UFMylation, respectively, and UFSP2, involved in pro-UFM1 biosynthesis and protein de-UFMylation, were found to be enriched in His_10_-UFM1ΔSC eluates by MS analysis, in line with their function during protein UFMylation (Figure 6C, left)^29,33^. UFM1 is covalently bound to UBA5 and UFC1 by thio-ester bond and transferred from UBA5 to UFC1 via a trans-thio-esterification reaction^29–31,80,82^. These proteins were also significantly increased upon induction of DSB by I-PpoI, suggesting an increase in nuclear/nucleolar UFMylation under these conditions (Figure 6C, right).

We furthermore identified several kinases known to be involved in DNA repair, including AURKA, AURKB and ATM as UFMylated proteins. While also detected in undamaged cells, UFM1-binding/UFMylation of ATM and AURKA was significantly increased upon DSB induction, in addition to AURKB and HAT1, a histone acetyltransferase which promotes insertion of histone H3.3 at DSBs (Figure 6C)^41,83,84^. Moreover, a number of protein essential for chromatin segregation and stability such as FANCD2, SMC6, PDS5A, and SMC2, as well as factors involved in DSB re-localization via the cytoskeleton and actin network (*i.e*., UNC45, SMC6) or regulating the actin network (*i.e*., ARHGAP11A, CDC42) were found to be UFMylated in response to DNA damage, several of which have known functions in, at least nuclear if not nucleolar, DNA damage repair^11,85–88^. Gene ontology analysis after rDNA DSB induction suggested that UFMylated proteins enriched upon I-PpoI induction are involved in chromosome organization (*e.g*., FANCD2, SMC6, ATM) and nuclear chromosome segregation (*e.g*., ASPM, AURKA/B, SMC2), consistent with the notion that UFMylation may be important for early steps of rDNA DSBs repair (Figure 6D-E).

Since induction of I-PpoI induces DSB both on genomic and ribosomal DNA, we cannot exclude that the observed UFMylation of known DNA repair factors and chromatin regulators occurs in response to genomic DSBs. However, I-PpoI has eleven genomic cleavage sites, compared with one within the 28S rRNA gene, the latter of which being present in ∼ 300 copies per cell. The high levels of nuclear cap formation observed upon I-PpoI induction suggest cleavage of I-PpoI in at least two-thirds of rDNA copies, which would represent an ∼18-fold excess of rDNA DSBs, suggesting that the majority of enrichment in protein UFMylation of repair factors is likely linked to rDNA DSB repair. Moreover, the identification of UFMylation of proteins with known roles in chromatin segregation, cohesion, and stability upon I-PpoI induction is in line with the required segregation of rDNA into nucleolar caps for subsequent repair. Lastly, we also observed co-purification of TCOF1 with His_10_-UFM1ΔSC; however, it was below our data cut off (data not shown).

Taken together, our data represents the first analysis of nuclear and nucleolar UFMylation targets in response to induction of DSBs. Moreover, based on the observation that predominately proteins with roles in DNA damage sensing (*e.g*., ATM, AURKA/B), chromatin regulation (*e.g*., HAT1) and segregation (*e.g*., ASPM, AURKA/B, SMC2, SMC6, UNC45), and initiating DNA repair (*e.g*., ATM) were found UFMylated upon induction of DNA damage, the results suggest that UFMylation may function as a regulator of early steps in the rDNA response pathway.

## DISCUSSION

Ribosomal DNA loci represent a unique region of the genome that is essential for cell viability. However, with its repetitive sequence and high transcriptional levels it also particularly prone to DNA damage and potential rearrangements during repair^89^. Therefore, distinctly regulated repair mechanisms, including nucleolar segregation of rDNA, are needed in order to ensure its accurate repair and maintain rDNA repeat stability. While some insights into nucleolar cap formation upon induction of rDNA damage have been made in recent years and current evidence suggests that silencing of rDNA transcription may be a pre-requisite for nucleolar cap formation^11^, there is still little understanding of how nucleolar cap formation, and the early stages of ribosomal DNA damage response are regulated.

### UFMylation as essential regulator of ribosomal DNA damage response

Recent work has identified that MRE11 and histone H4 are UFMylated in an early response to damage on nuclear DNA^34,35,37^. The MRE11-RAD50-NBS1 complex functions as a sensor of DNA DSBs, whereby MRE11-RAD50 bind the broken DSB ends and NBS1, in turn, recruits monomeric ATM to break sites, where the latter is activated through autophosphorylation^90,91^. UFMylation of MRE11 was suggested to potentially promote MRN complex formation and its recruitment to DSBs and the subsequent optimal activation of ATM, which was abrogated in the absence of the E3 UFM1-ligase UFL1^35,37^. The MRN complex is also recruited to damaged ribosomal DNA in a step that is instrumental for robust ATM and subsequent ATR activation; however, these steps are TCOF1-dependent and take place only upon nucleolar cap formation^12,14–16^. ATM was shown to be required for repression of ribosomal RNA synthesis upon induction of DNA double-strand breaks in rDNA repeats, an event suggested to precede the segregation of rDNA into nucleolar caps^70^, yet the signalling events that regulate these responses are largely elusive.

The identification of all components of the protein UFMylation pathway as sensitizers of ribosomal DNA damage as observed in our genetic screen, implicates UFMylation as an essential regulator in the rDNA damage response (Figure 1D; Table S2). Moreover, the fact that known downstream DNA damage response factors (*e.g*., BRCA2, 53BP1, and TOPBP1) were sensitizers (NormZ ≤ -0.17, - 0.73, and -1.11, respectively) but not identified as essential for rDNA DSB repair, suggests that the events prior to their recruitment – *i.e*., rDNA transcriptional silencing and nucleolar segregation – are crucial for rDNA DSB repair to ensue and that UFMylation may be required for the regulation of these early processes on damaged rDNA. This is further supported by the identification of ATM as UFMylation target and the significant enrichment of its UFMylated form upon I-PpoI induction (Figure 6C, right; Table S9). Previous work demonstrated that rDNA transcriptional silencing upon I-PpoI rDNA DSB induction is ATM-dependent and induces nucleolar reorganization at the nucleolar periphery ^10,13,16,38,70^. The observation that cells lacking the E3 UFM1-ligase UFL1 exhibit reduced rDNA transcriptional silencing and overall reduced nucleolar segregation upon induction of rDNA DSB (Figure 3A-E) suggests that UFMylation of ATM is likely essential for these steps to be carried out efficiently^34–36^, and that UFMylation of ATM may be part of previously described positive feedback loops whereby ATM phosphorylation of UFL1 at Ser462 enhances UFL1 activity, resulting in increased ATM UFMylation and activity in rDNA silencing (Figure 7)^34–36^. Moreover, the reduced levels of Ser1981-phosphorylated ATM in absence of UFL1, suggest that ATM UFMylation may take place prior to its phosphorylation at Ser1981, or that ATM UFMylation may also affect efficient ATM autophosphorylation at this site post-rDNA silencing and prior to nucleolar segregation ^34,35,37^.

**Figure 7.**
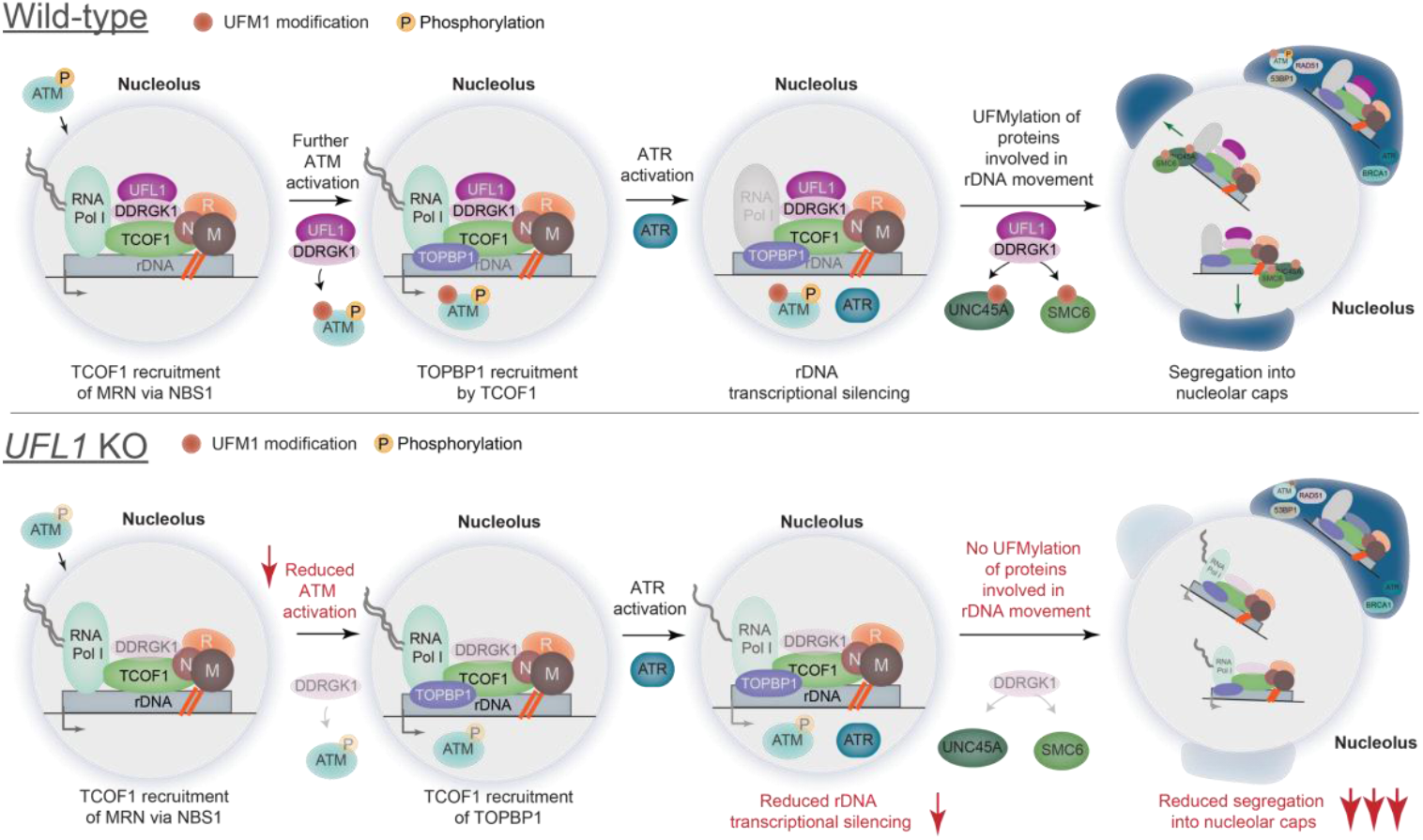
Model of UFMylation in rDNA DSB repair. Upon induction of rDNA DSBs, initial activation of ATM allows for MRN recruitment to nucleoli via the TCOF1-NBS1 interaction. UFL1 and DDRGK1 are also recruited to nucleoli to activate ATM, potentially through direct UFMylation of ATM. Nucleolar MRN and ATM activation allows for TOPBP1 recruitment and ATR activation, allowing for rRNA transcriptional silencing, nucleolar segregation into caps and downstream DNA repair protein recruitment. As UFL1 and DDRGK1 are associated with TCOF1 within nucleolar caps, they may also potentially UFMylate proteins that may mediate either rRNA transcriptional silencing or rDNA DSB movement to the nucleolar periphery, such as UNC45A and SMC6. UFL1-deficient cells have reduced protein levels of DDRGK1, impaired ATM activation, reduced rRNA transcriptional silencing, and an overall reduction in nucleolar segregation.

The observation that UFL1 and DDRGK1, which stabilizes UFL1 and enhances its activity, localize to nucleolar caps upon rDNA damage and are isolated with TCOF1-associated complexes (Figure 5A, C) further supports a role for UFMylation in the maintenance and activation of rDNA damage response^13,42,78^. The partial loss of rDNA silencing and nucleolar segregation in the absence of UFL1 raises the question whether UFMylation is specifically required for increased ATM activation and rDNA silencing under persistent rDNA damage conditions such as I-PpoI induced damage^10^. Alternatively, DDRGK1 may have the potential to direct UFMylation of so far unidentified E3 UFM1-ligases. UFMylation of several substrates by UFL1 was shown to rely on DDRGK1 (*e.g*., RAB1A/B, RAB5C, ARF4, clathrin)^29^. The observation that both UFL1 and DDRGK1 can exist in the nucleus independently of one another (Figure S5), suggests that potential alternative binding partners may exist in the nucleus or nucleolus that can confer alternative substrate specificity, even if to a lesser extent, as loss of UFL1 results in a less efficient rDNA silencing and damage response.

It is also still unknown if UFL1 and DDRGK1 interact with TCOF1 directly or if they are associated with TCOF1 through the MRN complex, as UFL1 was previously found to bind to the MRN complex in a DNA damage-dependent manner and UFMylate MRE11^37^. We isolated UFMylated MRE11 with His_10_-UFM1 upon I-PpoI induction, but, similar to Histon H4 whose UFMylation was suggested to regulate, at least in part, ATM activation on nuclear DNA either through affecting H3K9 methylation or changes in local chromatin structure^34^, it was not significantly enriched over background controls, suggesting either mono-UFMylation or that MRE11 and H4 UFMylation occurs predominantly at nuclear DNA DSB sites, which are underrepresented in our system^29^. Similarly, while UFMylated TCOF1 was purified with His_10_-UFM1 upon damage, it was not significantly enriched. As such the UFMylation status of TCOF1 and its subsequent impact on rDNA damage repair remain to be determined.

### UFMylation mediates rDNA chromatin reorganization and nucleolar segregation

Previous data suggest that ATM activation following rDNA DSB precedes nucleolar segregation^10,14^. However, activation of ATM requires changes in chromatin such as histone modifications and opening of chromatin that are critical for ATM activation, and H3K9 and H3K36 trimethylation as well as H4K16 acetylation were all suggested to regulate ATM activity^10,92^. Recent data showed the downstream effector human silencing hub (HUSH) complex, comprised of TASOR, MPP8, and perihpillin, to mediate H3K9Me3 deposition on damaged ribosomal DNA leading to heterochromatin formation and transcriptional silencing^10–13^. The identification of the HUSH component *FAM208A* (encoding TASOR) and the effector H3K9Me3 methyltransferase *SETDB1* as essential for cell viability in response to rDNA DSBs further highlights the crucial importance of an effective early rDNA damage response for cell survival (Figure 1G; Table S2),^11^ but is also in contrast to previous findings that SUV39H1/2, rather than SETDB1, was important for H3K9Me3 deposition at the rDNA locus^11^. This observed difference may be due to a difference in persistence of rDNA DSBs induction as AsiS-I, in contrast to I-PpoI, does not generate DSBs at all rDNA repeats^11^. Since I-PpoI generates a greater amount of rDNA DSBs, there may be an increased requirement for rDNA transcriptional silencing, which may involve additional mechanisms unrelated to H3K9-trimethylation, such as SETDB1-mediated methylation of UBF ^93^.

SUV39H1/2, however, has also been linked to rDNA heterochromatin maintenance in connection with the known rDNA chromatin remodeler <u>a</u>lpha-thalassemia mental retardation <u>X</u>-linked (ATRX) through SUV39H1/2-mediated K9Me3 of histone H3.3^53,54^. H3.3 is enriched at both actively transcribed as well as several transcriptionally silent regions including telomeres and pericentric heterochromatin^53,57,94–97^. Moreover, H3.3 was also shown to be rapidly deposited at sites of UV damage in nuclear DNA and suggested to facilitate recovery of transcription post-DNA repair^98^. H3.3 deposition is, in part, mediated by the chromatin remodeler ATRX, which together with its chaperone DAXX incorporates H3.3 in pericentromeric and telomeric regions but was also found to regulate DNA repair during homologous recombination^55,56^. In mouse ES cells, ATRX-mediated H3.3 deposition was also demonstrated to contribute to rDNA heterochromatin formation and stability, as loss of ATRX led to reduced levels of H3.3, H3K9me3, and H4K20me3, increased aberrant recombination events of rDNA repeats, increased nucleolar fragmentation and formation of extrachromosomal rDNA circles^54^. The identification of *ATRX* as a gene that sensitizes cells to rDNA damage supports its importance in mediating rDNA heterochromatin formation upon rDNA damage (Figure 1D; Table S2), in line with a role for ATRX in the maintenance of rDNA loci stability, beyond mouse ES cells. Beside ATRX, H3.3 deposition is also mediated by the <u>hi</u>stone regulator <u>A</u> (HIRA). The HIRA-CABIN1-UBN1 complex deposits H3.3-H4 predominantly at actively transcribed genes, and controls chromatin integrity and H3.3 occupancy at these regions^57–61^. Notably, HIRA has been implicated in transcriptional restart after UV-induced DNA damage via deposition of H3.3 into chromatin around UV lesions as well as rDNA transcription in mouse zygotes^61,99^. The observation that *HIRA, CABIN*, and *UBN1* are cell sensitizers to rDNA damage (Figure 1F; Table S2) implicates them for the first time in rDNA repair, further underscoring the importance of maintaining faithful rDNA repeat integrity, while coordinating rDNA repair and post-repair transcriptional restart^85^.

Our observations also point towards an important role for H3.3 in rDNA repair; first, in the establishment and maintenance of heterochromatin for rDNA silencing as prerequisite for nucleolar segregation and repair, and second, in facilitating HIRA-mediated transcriptional restart. Moreover, in line with UFMylation as regulator of critical early steps necessary for rDNA silencing and nucleolar segregation, we observed UFMylation of the histone acetyltransferase 1 HAT1 in response to I-PpoI induction (Figure 6C, right; Table S9). In both *S. cerevisiae* and mammalian cells, HAT1 is required for the promotion of efficient DNA DSB repair by homologous recombination, in the latter via the incorporation of H4K5/K12-acetylated H3.3 at sites of double-strand breaks mediated by HIRA to facilitate subsequent recruitment of key repair factors^83,100^, again linking regulation of early chromatin changes to efficient repair.

While DSB segregation and nucleolar cap formation has previously been proposed as a direct consequence of DSB-induced rDNA transcriptional silencing^13^, more recent work suggests that nucleolar segregation may instead be an active transcription-independent process, mediated by the actin network and the nuclear envelope-associated ‘<u>Li</u>nker of <u>N</u>ucleoskeleton and <u>C</u>ytoskeleton’ (LINC) complex^11^. The LINC complex, ARP3, a component of the ARP2/3 and regulator of the actin cytoskeleton, and UNC45A, a myosin regulator, were demonstrated to mediate rDNA DSB mobilization at the nucleolar periphery as depletion of all three factors (SUN1, UNC-45A, and ARP3) did affect rDNA segregation but not rDNA transcriptional repression following DSB induction^11^. Collectively, these data suggest that the nuclear envelop, LINC complex and components of the actin network ensure rDNA DSB mobilization downstream from transcriptional inhibition^11^.

Notably, purification of nuclear and nucleolar UFMylated proteins after rDNA DSB induction, identified several proteins involved in cell cycle control as well as chromosome organization and segregation. Moreover, we observed UFMylation of UNC45A as well as I-PpoI damage-dependent UFMylation of SMC6 (Figure 6C; Table S9). UNC45A, a myosin activator, was previously reported to be important not only for movement of heterochromatic DSBs, but also for rDNA re-localization into segregated nucleolar caps^11,85^. SMC6 is part of the SMC5/6 complex and essential for accurate rDNA segregation and recombination-mediated repair in yeast and human cells^5,101–105^. SMC5/6 is highly enriched with rDNA, where its associated E3-SUMO ligase NSE2/MMS21 is important for nucleolar heterochromatin DSB re-localization^85–87^. Our observations are in line with a role for the actin network in nucleolar segregation and a role for UFMylation as key regulator of early rDNA events including nucleolar cap formation. In such a model, UFMylation of ATM and HAT1 would not only mediate chromatin changes and rDNA silencing but UFMylation of components if the actin and myosin network would further regulate either their localization to or function in nucleoli, enabling and actively mediating re-localization of rDNA DSBs to the nucleolar periphery (Figure 7).

### Final Remarks

Although the precise mechanisms of UFMylation in response to rDNA DSBs remain to be determined, our work provides important insights into the current model of TCOF1-dependent rDNA repair. Given our observation that all components of the UFMylation pathway are essential for rDNA repair and the identification of UFMylated key factors implicated in the rDNA damage response such as ATM and HAT1 as well as several actin and myosin network components upon induction of rDNA DSBs suggest that UFMylation regulates in particular early steps of the rDNA damage response, including ATM activation, leading up to nucleolar segregation and rDNA silencing. Interestingly, the observations that FANCI and FANCD2 are UFMylated upon I-PpoI induction also suggest the potential activation of multiple repair pathways. Indeed, recent work suggested a role for Fanconi anemia pathway in the repair of CRISPR/Cas9-induced DSBs or persistent breaks, which has yet to be determined^106,107^, unless UFMylation can also function as inhibitor, a potential function that has not yet been investigated. Overall, the identification of UFMylation as a crucial regulator of rDNA DSB repair and key UFMylation targets in response to rDNA DSBs provides new insights in the early steps of ribosomal DNA repair, and future studies will further elucidate how exactly UFMylation regulates these pathways, whether through mediation of protein-protein interactions and recruitment or via direct modulation of protein activity.

## DATA AVAILABILITY

New data reported in this article are available in the online supplementary material. MAGeCK sgRNA read counts are reported in Supplemental Table 1 and normalized mass spectrometry data in Tables S8 and S9. Mass spectrometry data has been deposited to the ProteomeXchange Consortium via the PRIDE partner repository (Project ID PXD048568). All raw data and analysis pipelines will be shared on request to the corresponding author.

## SUPPLEMENTARY DATA

Supplementary Tables are available <u>online</u>.

## ACKNOWLEDGEMENTS

We would like to acknowledge the IRCM core facilities - Genomics (S. Boissel), Proteomics (D. Faubert), Flow Cytometry (J. Lord), Microscopy (D. Fillion) - for their support. Dr. A. Orthwein for the hTERT-RPE1 WT and hTERT-RPE1 *p53-/-* Cas9 cell lines. We thank L.C. Aguilar, Dr. N. Francis, T. Ubhi, S. Mockly, S. Ehresmann, E. Goretti and J.F. Laurendeau for feedback and critical reading of the manuscript.

## AUTHOR CONTRIBUTIONS

P.P. and M.O. conceived the study. P.P., E.D., M.G., and LC.A. performed all experiments. C.P. performed data analysis. M.O. and J.V. supervised the work, and P.P. and M.O. wrote the paper with input from all authors.

## MATERIAL AND METHODS

### Cell lines and cell culture conditions

All human cell lines used in this study are listed in Table S6. hTERT-RPE1 WT and hTERT-RPE1 *p53-/-* Cas9 cells were a gift from Alexandre Orthwein. Cells were cultured at 37°C in 5 % CO_2_ in Dulbecco’s modified Eagle’s medium (DMEM) supplemented with 1 % PenStrep and 10 % fetal bovine serum (FBS, Wisent). hTERT-RPE1 cells expressing TCOF1-eGFP and eGFP-NLS were generated by integrating pSH-EFIRES-TCOF1-eGFP into the AAVS1 safe-harbor locus by co-transfection with pCas9-sgAAVS1-#1/#2 using lipofectamine 3000 as previously described^108^. Cells were trypsinized 72 hr post transfection and resuspended in Hanks Balanced Salt Solution (HBSS) + 2% FBS. GFP positive cells were then bulk sorted on a BDFACS AriaIII cell sorter. hTERT-RPE1 *p53-/-* Cas9 expressing DD-HA-ER-I-PpoI were generated by lentiviral integration of pLX303-DD-HA-ER-I-PpoI -G418 at an MOI <1 and selected using 800 μg/mL G418. The same cells expressing 10xHis-UFM1ΔSC were selected using GFP positive bulk sorting as described below.

### CRISPR-Cas9 gene editing and single cell cloning

sgRNAs targeting UFL1, DDRGK1, or LacZ were cloned into Lenti-Guide-Puro vectors using Zhang lab protocol^109^. After selection with puromycin, cells were trypsinized, resuspended in 1 X HBSS + 2 % FBS and sorted into single cells in a 96-well plate using a BD FACS AriaIII. Alternatively, single cells were plated at an average of 0.5 cells per well in 96-well plates by serial and limiting dilution. Individual clones were validated by western blotting and Sanger sequencing. sgRNA sequences are listed in Table S7.

### Plasmids and viral vectors

DD-HA-ER-I-PpoI (a gift from Michael Kastan, Addgene: 49052) was subcloned into a pLX303 backbone vector (a gift from Mikko Taipale, Addgene: 154472) using In-Fusion cloning. The G418 (Geneticin) resistance gene was subcloned from pDEST-CMV-C-eGFP (a gift from Robin Ketteler, Addgene: 122844) and inserted into pLX303-DD-HA-ER-I-PpoI using In-Fusion cloning. eGFP vectors were cloned using Gateway cloning from pDONR221-eGFP (a gift from David Root, Addgene plasmid # 25899) and pFRT-TO-DEST (a gift from Dr. Thomas Tuschl, Addgene plasmid # 106348). An NLS was then added to eGFP using standard PCR. TCOF1 was sub-cloned into pFRT-TO-DEST using In-Fusion cloning. C-terminal eGFP was cloned using Gateway cloning. TCOF1-eGFP was then cloned into pSH-EF-IRES-AtAFB2-mCherry (a gift from Elina Ikonen, Addgene plasmid # 129716) using In-Fusion cloning. eGFP-NLS was cloned in a similar manner. sgRNAs were cloned into LentiGuide-Puro (a gift from Feng Zhang, Addgene: 52963) or a modified form of LentiGuide-Puro in which Cas9 was replaced by GFP-NLS or mCherry-NLS as previously described^110^. 10xHis-UFM1ΔSC was subcloned from pRK5-HA-UFM1 (a gift from Yihong Ye, Addgene: 134639) using PCR and inserted into a modified Lenti-Guide-Puro GFP by replacing the puromycin selection marker with 10xHis-UFM1ΔSC using In-Fusion cloning. A list of all plasmids used in this paper is provided in Table S11.

### Immunofluorescence microscopy

For immunofluorescence microscopy experiments, 3.0×10^5^ cells were seeded onto 18-mm coverslips in 6-well tissue culture plates overnight and treated as described above. Cells were washed with PBS and fixed in 3.7% paraformaldehyde/1X PBS for 10 min at room temperature. Cells were then washed three times in 1XPBS for 5 min, followed by permeabilization for 10 min at room temperature in 0.3% Triton X-100/1XPBS. Cells were then washed three times in PBS for 5 min and blocked for 1 hr in blocking buffer (10% goat serum in 1X PBS). Samples were incubated overnight at 4°C diluted in blocking buffer in a humidified chamber. Coverslips were then washed three times in 1XPBS for 5 min, followed by incubation in secondary antibodies for 1 hr at room temperature. Finally, coverslips were washed 3 times in 1XPBS for 5 min and mounted using Prolong Gold containing DAPI. For immunofluorescence of endogenous UFL1, cells were fixed in 3.7% paraformaldehyde for 10 min on ice and permeabilized with 0.5% Triton X-100 for 10 min at room temperature. Primary antibody against UFL1 was incubated for 1 hr at room temperature. Imaging for biological replicates was carried out under identical conditions. Antibodies and antibody dilutions are provided in Table S10.

### Image acquisition

Images were acquired on a confocal microscope (SP8, Leica) using 63x objective at 2,048 × 2,048 pixels per image. Images were acquired in sequential line-scan mode. For Z-stacks, a minimum of 6 slices were taken at a system optimized distance of at least 0.33 μM.

### Image quantification

Images were exported using LASX and processed using Fiji. Co-localization analysis and quantification were performed as previously described^111^. Briefly, single cells were isolated from single-slice z-stack images and imported into Fiji. ComDetv0.5.2 was then used to quantify total foci in each channel as well as co-localized foci between channels. ComDet detection settings are described in Table S5. For determination of cells with fully segregated nucleolar caps, images were imported into Fiji and quantified manually. Line scans for co-localization were performed using the ‘Plot Profile’ function in Fiji.

### Clonogenic survival assays

hTERT-RPE1 *p53-/-* Cas9 expressing DD-HA-ER-I-PpoI and indicated sgRNAs were seeded in 6-well plates (400 cells per well) and treated with 100 nM Shield-1 and 100 nM 4-OHT 1 day after plating and every 3 days thereafter. After 10-14 days, colonies were stained with crystal violet (0.4 % w/v in 20% methanol) and manually counted. Relative survival was calculated as previously described^112^. Data is represented as a ratio of cells treated with Shield-1+4-OHT divided by DMSO per biological replicate.

### Protein extraction and immunoblotting

Cells or nuclei were lysed in 3:1 volume to pellet volume in RIPA buffer (50 mM Tris-HCl pH 8.0, 150 mM NaCl, 1 % NP-40, 0.5 % sodium deoxycholate, 0.1 % SDS, 1 mM MgCl_2_) containing benzonase (Sigma) for 30 minutes on ice before centrifugation at 21,100 x g for 10 min at 4°C. Protein concentration was quantified using the Lowry DC Assay (Bio-Rad). Proteins were separated on a 6 %, 10 % or 4-15 % Tris-Glycine SDS-PAGE gels and transferred to either 0.2 mM nitrocellulose or 0.2 mM PVDF membranes. Alternatively, 4-12% Bis-Tris NuPage gels were used. Membranes were blocked for 1 hr at RT using 5% skim milk powder in PBS-T prior to detection with antibodies as indicated in Table S10. Proteins were detected using the Licor Odyssey system.

Signal intensities were quantified using Licor Imagestudio Lite. Band intensities for ATM and pATM-S1981 were first normalized against GAPDH. The ratio of ATM to pATM-S1981 was then calculated and expressed as a ratio over their levels in DMSO treated cells.

### Propidium Iodide staining and cell cycle analyses

Cells were collected after treatment by centrifugation at 1000 x g for 5 min at room temperature, washed once in PBS, resuspended in HBSS containing 2% FBS and then fixed using ice-cold 70 % ethanol over a low-speed vortex. Fixed cells were washed twice with HBSS containing 2 % FBS before resuspension in HBSS containing 0.1 mg/mL propidium iodide (Sigma), 2 mg/mL RNAseA (Thermo), and 0.6 % NP-40. Cells were transferred to a 5-mL round-bottom polystyrene tube and incubated at room temperature for 30 min prior to filtering through a 40 μm cell strainer. Data was collected on the FACSCalibur using the FL3 (670LP) filter with a minimum of 20000 events. Data analysis was carried out in FlowJo vX0.7. Cell counts are displayed using the modal option.

### CRISPR-Cas9 genetic screening and DNA Purification

hTERT-RPE1 *p53-/-* Cas9 expressing DD-HA-ER-I-PpoI were transduced with the lentiviral TKOv3 library at a low MOI (∼0.30) in the presence of 8 μg/mL polybrene (Sigma)^43,44^. TKOv3 lentivirus was produced as previously described^113^. Transduced cells were selected using puromycin (20 μg/mL) as previously described^72^. The initial time 0 (T0) timepoint was 3 days after initial infection, in which cells were pooled together and split into 2 technical replicates “A” and “B”. Pooled cells were then expanded for 9 days (passaging every 3 days) after which they were split into 2 groups, DMSO vehicle control treated cells as well as cells that were treated with 100 nM Shield-1 (Medchemexpress) and 100 nM 4-hydroxytamoxifen (4-OHT, Medchemexpress). The concentration of Shield-1 and 4-OHT used to treat cells were determined using LD20 curves determined over 20 days (Figure S1E). Cells were treated for 1 hr at 37°C 1 day after plating, washed twice with PBS and replenished with fresh media. Cells were passaged and treated every 3 days until the end of the experiment at timepoint 24 (T24). Cell pellets were collected in technical replicates, snap frozen, and stored at -80°C. Genomic DNA corresponding to Timepoint 16 (T16) and 24 (T24) was extracted using the Wizard Genomic DNA Purification kit as per manufacturers protocol and resuspended in 400 μL of TE pH 8.0.

### Library preparation and next generation sequencing

The libraries were prepared with a two-step PCR to (1) enrich guide-RNA regions in the genome and (2) amplify guide-RNAs with Illumina TruSeq adapters with i5 and i7 indices. PCR 1 was performed on 140 μg to 200 μg of genomic DNA (42 PCR reactions per sample, 25 cycles of amplification) using the LCV2 forward and reverse primers. All PCR products of the same sample were pooled, purified with KAPA Ampure beads (1X) and size-selected with KAPA Ampure beads (L:1 / R:0.5). PCR Reverse primers (10 cycles) and purified with KAPA Ampure beads (1.2X). Size selection (range 100 bp-190 bp) was carried out using a Pippin Prep 2% agarose cassette followed by a purification with KAPA Ampure beads (1.2X). Size distribution of the final libraries was assessed on a Bioanalyzer hsDNA chip and libraries were quantified by qPCR. Libraries were sequenced on an Illumina NextSeq 500 High Output v2.5 75 Cycles flowcell with the following parameters: Read 1: 21 (Dark Cycles) + 26, Index Read 1: 8, Index Read 2: 8 Primer sequences have been previously described in^43,65^ and are listed in Table S4.

### Data processing, quality control and gene essentiality scoring

The overall quality of data (mapped reads, Gini index) was first assessed using MAGeCK^114^. The sgRNA read count files were calculated using the raw fastq files for each experiment using the MAGeCK ‘count’ function on Galaxy Australia. Secondly, BAGELv2 was used to calculate precision-recall curves using the ‘pr’ function using the core essential and nonessential gene lists^45^. Log_10_ sgRNA counts for each sgRNA corresponding to core essential and nonessential genes were plotted for each replicate and timepoint as a readout of screen quality. MAGeCK computed read count files were then used to calculate normZ gene scores using the DrugZ algorithm comparing I-PpoI induced replicates to their uninduced replicates at each paired timepoint. Aligned sgRNA read counts are provided in Table S1, with DrugZ analyzed datasets for T16 and T24 provided in Tables S2 and S3. GO:Term enrichment was performed using g:Profiler^46,115^.

### Competitive growth assay

Immediately after selection, 5×10^4^ of the hTERT-RPE1 cells transduced with UFL1 sgRNAs (in Lentiguide-gRNA-NLS-GFP-2A-PURO) were seeded with 5×10^4^ of cells transduced with an AAVS1 targeting sgRNA (in Lentiguide-NLS-mCherry-2A-PURO) in 12-well plates and incubated overnight. The following day, the plates were imaged using the 4x objective of IncuCyte (Sartorius). To assess the representation of the green and red populations, the fluorescent intensity of GFP as compared to mCherry was measured every 24 hr and normalized to Day 0 of the assay. Cells were passaged using a 1:2 dilution every 3 days. 1 Day after passaging, cells were treated with 100 nM Shield-1+ 100nM 4-OHT to recapitulate conditions from the CRISPR-Cas9 genetic screen. In total, cells were grown and passaged for 21 days.

### Cryogenic GFP-affinity purification

5.0×10^6^ hTERT-RPE1 cells expressing TCOF1-eGFP were plated into 15cm tissue culture dishes and allowed to grow overnight. The following day, cells were treated with 2 μM Shield-1 + 2 μM 4-OHT for 6 hr at 37°C. Cells were scraped and collected in PBS and centrifuged at 1000xg for 5 min at 4°C. The cell pellets were transferred to a pre-weighed 15mL tube and centrifuged at 1000xg for 5 min at 4°C. Cryogenic milling was carried out as previously described^116^, with the following modifications. Briefly, excess PBS was removed, and the weight of the wet cell pellet was measured using an analytical scale. Cells were resuspended in 1:1 weight to volume (1 μL per 1 mg) lysis buffer (50 mM Tris-HCl pH 8.0, 100 mM NaCl, 10% glycerol, 0.5% Triton X-100, 0.01% NP-40 containing 1 X protease inhibitor cocktail (Sigma), 5 mM NaF, 1 mM NaPyPh, 1 mM BGP, 1 mM NaO). Cell droplets were created by pipetting the resuspended cells into liquid nitrogen. ∼100 mg of cell droplets was then transferred to 2 mL screw cap tubes containing 3×5 mm stainless steel metal balls and lysed using a Retsch MM400 for 3 min at 25 Hz to create a cell powder. The cell powder was resuspended in a 1:1 weight to volume ratio (1 μL per 1 mg) in lysis buffer. Where indicated, nucleic acid digestion was performed by addition of Benzonase at 1:1000 and supplemented with 1 mM MgCl_2_ and incubated on ice for 30 min. Extracts were cleared by centrifugation at 21,100xg for 10 min at 4°C. Protein concentration was quantified using the Lowry DC assay (Bio-Rad). 10 mg of quantified lysates were incubated with 15 mg of GFP-nanobody conjugated magnetic beads (Dynabeads M-280) for 1 hr at 4°C on a nutator. GFP-nanobody was expressed from pDZ580-pET28a-GBP and purified as previously described^117^. After removal of the supernatant, beads were washed with lysis buffer, followed by 100 mM NH_4_OAc + 1 mM MgCl_2_ to remove detergents before a final wash and resuspension in 20 mM Tris-HCl pH 8.0. Purified protein complexes were digested on-bead at 37°C with 500 ng trypsin for 16 hr at 350 RPM in a thermomixer. Trypsin digestion was stopped by addition of formic acid to 2%. Peptides were then dried in a speedvac.

### Subcellular fractionation and nuclear extraction

2.5×10^6^ hTERT-RPE1 cells expressing were plated into 10 cm tissue culture dishes and allowed to grow overnight. The following day, cells were treated with 2 μM Shield-1 + 2 μM 4-OHT or DMSO for 6 hr at 37°C. Cells were scraped and collected in PBS and centrifuged at 1000xg for 5 minutes at 4°C. Cells were first lysed with hypotonic lysis buffer (50 mM Tris-HCl pH 7.5, 10 mM NaCl, 1.5 mM MgCl_2_, 10 % glycerol, 0.34 M sucrose) in a volume 10X the volume of the cell pellet. Lysed cells were incubated on a nutator for 10 min at 4°C, incubated on ice for 10 min followed by addition of Triton X-100 to 0.1%. Nuclei were sedimented by low-speed centrifugation at 1,300 x g for 10 min at 4°C. The remaining supernatant was stored as the cytoplasmic fraction. Nuclei were washed once more with hypotonic lysis buffer and stored at -80°C.

### His_10_-UFM1ΔSC affinity purification

5.0×10^6^ hTERT-RPE1 cells expressing 10xHis-UFM1 were plated into 15 cm tissue culture dishes and allowed to grow overnight. The following day, cells were treated with 2 μM Shield-1 + 2 μM 4-OHT or DMSO for 6 hr at 37°C. Nuclei were purified as described in *subcellular fractionation and nuclear extraction*. Purified nuclei were lysed in denaturing urea buffer (8 M urea, 100 mM Na_2_HPO_4_/NaH_2_PO_4_, 300 mM NaCl, 10 mM Tris-HCl pH 8.0, 10 mM imidazole pH 8.0). Lysed nuclei were sonicated using a Branson sonicator (3 cycles of 30 % amplitude, 10 sec) and cleared by centrifugation at 21,100 x g for 10 mins at RT. Protein concentrations were quantified using the Lowry DC assay (Bio-Rad). 2.5 mg of nuclear extracts were bound to 50 μL of magnetic HisPur Ni-NTA beads (ThermoFisher Scientific) for 1 hr at RT on a rotating wheel. Beads were washed 8 × 1 mL with wash buffer (8 M urea, 100 mM Na_2_HPO_4_/NaH_2_PO_4_, 300 mM NaCl, 10 mM Tris-HCl pH 8.0, 20 mM imidazole pH 8.0), 1 x with wash buffer without urea and 3 x with 1 mL of 50 mM ammonium bicarbonate before on-bead digestion using 500 ng of trypsin for 16 hr at 37°C in 50 mM ammonium bicarbonate. Trypsin digestion was stopped by addition of formic acid to 2%. Peptides were then dried in a speedvac.

### Northern blotting

3×10^6^ cells were seeded into 10 cm tissue culture dishes and incubated overnight. After treatment as indicated in text, total RNA was extracted using TRIzol (Invitrogen) as per manufacturer’s instructions. 3-5ug of total RNA was separated on 1% agarose-formaldehyde gels in Tricine-Triethanoloamine gels as previously described^118^. RNA was transferred to a nylon membrane overnight by capillary action in 10X SSC, crosslinked using 1200 J of 254 nm short wave UV and incubated with probes labelled with DyLight 800 NHS Ester^119^. Probes were incubated overnight at 42°C in hybridization buffer (6X SSPE, 5X Denhardt’s, 0.2 mg/mL ssDNA, 0.2% SDS) with gentle agitation. rRNA precursors were detected using probes hybridizing with the 5’ ETS (5’-CGCTAGAGAAGGCTTTTCTC-3’). Membranes were washed twice with 2X SSC/0.1% SDS at 45°C, followed by one wash of 1X SSC/ 0.1% SDS at 45°C. 45/47S rRNA visualized using a Licor Odyssey system. Mature 28S and 18S rRNA visualized by methylene blue staining. Quantification was carried out using ImageStudio Lite.

### EU transcription assay

3×10^5^ cells hTERT-RPE1 WT or *UFL1* KO cells were seeded onto 18 mm coverslips and allowed to grow overnight. The following day, cells were treated with 2 μM Shield-1 + 2 μM 4-OHT or DMSO for 5 hr at 37°C. Cells were then incubated with 1 mM EU for one additional hour (6hr total incubation time) before fixing with 3.7 % w/v paraformaldehyde and permeabilized with 0.5 % Triton-X-100 in PBS. Click-IT Alexa594 (Life Technologies) reactions were performed as per manufacturer’s instructions. After Click-IT reactions, cells were labelled with TCOF1 as previously described in the immunofluorescence section. Quantification of EU signal per nucleus was measured using CellProfiler. Briefly, maximum intensity projections were made using the Leica LASX software and imported into CellProfiler. TCOF1 foci were detected and segmented such that only EU signal that overlapped with TCOF1 signal was quantified. The total EU signal per nucleus was summed in R-Studio and used for downstream quantification. Only cells with nucleolar EU signal (A.U.) >10 were quantified.

### LC-MS/MS

Desalting/cleanup of the digests was performed using C18 ZipTip pipette tips (Millipore, Billerica, MA). Eluates were dried down in vacuum centrifuge and stored at -20ºC until LC-MS/MS analysis. Samples were reconstituted under agitation for 15 min in 16.5 μL of 2% acetonitrile -1% formic acid and loaded into a 75 μm i.d. × 150 mm Self-Pack C18 column installed in the Easy-nLC II system (Proxeon Biosystems). Peptides were eluted with a three-slope gradient at a flowrate of 250 nL/min. The buffers used for chromatography were 0.2% formic acid in water (solvent A) and 0.2 % formic acid in acetonitrile (solvent B). Solvent B first increased from 2 to 15 % in 30 min, from 15 to 36 % B in 60 min and then from 36 to 80 % B in 10 min. The HPLC system was coupled to Orbitrap Fusion mass spectrometer (Thermo Scientific) through a Nanospray Flex Ion Source. Nanospray and S-lens voltages were set to 1.3-1.7 kV and 60 V, respectively. Capillary temperature was set to 250 °C. Full scan MS survey spectra (m/z 360-1560) in profile mode were acquired in the Orbitrap with a resolution of 120,000 with a target value at 3e5. A top 3s method was used for the most intense peptide ions that were fragmented in the HCD collision cell and analyzed in the linear ion trap with a target value at 2e4 and a normalized collision energy at 29 V. Target ions selected for fragmentation were dynamically excluded for 15 sec after two MS2 events.

### Protein identification

The peak list files were generated with Proteome Discoverer (version 2.3) using the following parameters: minimum mass set to 500 Da, maximum mass set to 6000 Da, no grouping of MS/MS spectra, precursor charge set to auto, and minimum number of fragment ions set to 5. Protein database searching was performed with Mascot 2.6 (Matrix Science) against the UniProt Human protein database. The mass tolerances for precursor and fragment ions were set to 10 ppm and 0.6 Da, respectively. Trypsin was used as the enzyme allowing for up to 1 missed cleavage. Methionine oxidation was specified as a variable modification. Data interpretation was performed using Scaffold version 5.1.2.

### Mass Spectrometry Data Analysis

Protein and peptide identification thresholds in Scaffold were set to 95 % allowing for a false discovery rate of ∼5 %. Semi-quantification of proteins detected in TCOF1-eGFP and eGFP-NLS affinity purifications were carried out as previously described^120^. Briefly, only proteins which contained at least one Exclusive Unique Spectral Count (EUSC) were filtered and analyzed using Exclusive Spectral Counts (ESCs). Only ESCs detected in affinity purified samples (*e.g*., TCOF1-eGFP) with counts < 2 ESCs in negative controls (*e.g*., eGFP-NLS) were retained. *In silico* digestion was performed using MS digest (http://prospector.ucsf.edu) to control for protein size and predicted trypsin cleavage sites. ESCs were then normalized against the average values of TCOF1-eGFP for each individual replicate. Normalized spectral counts were compared between DMSO and Shield-1+4-OHT treated samples using Student’s t-test. Fold-change was determined by taking a ratio of normalized spectral counts between DMSO or Shield-1+4-OHT treated samples. For 10xHis-UFM1ΔSC analysis, protein and peptide identification thresholds in Scaffold were set to 95 % allowing for a false discovery rate of ∼5 % prior to export of iBAQ intensity values. iBAQ intensities for each protein were compared to wild-type cells and significance was determined using Student’s t-test. Fold-change was determined using iBAQ ratio of DMSO or Shield-1+4-OHT treated cells as compared to wild-type cells. GO:Term enrichment was performed using g:Profiler^115^. Normalized spectral counts for identified proteins from the TCOF1-eGFP affinity purifications are provided in Table S8. Raw iBAQ intensity values for identified UFMylated proteins are provided in Table S9. Data are available via ProteomeXchange with identifier PXD048568.

### Data visualization, quantification, and statistical analysis

All data reported are representative of three independent biological replicates. Data analysis was performed in R and Prism 8 (GraphPad Software). Data was visualized using ggplot2. Statistical tests are described in each figure legend respectively (test used, n-value for cells). An alpha value of p=0.05 was used for statistical significance.

---

**Figure S1.**
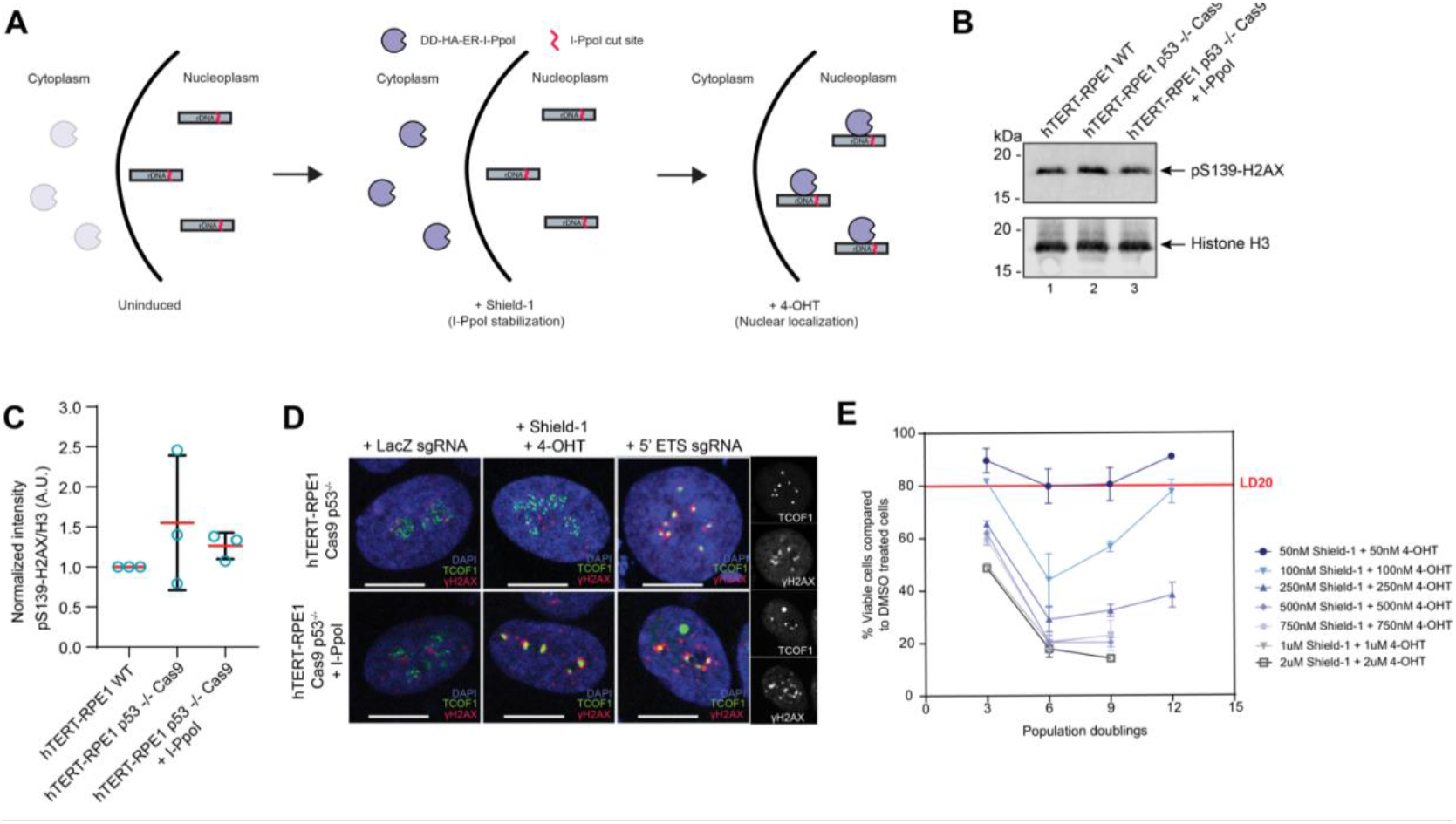
Characterization of hTERT-RPE1 *p53*^-/-^ Cas9 cells expressing DD-HA-ER-I-PpoI. **A)** Schematic of the DD-HA-ER-I-PpoI system. In uninduced (*e.g*., DMSO control) conditions, I-PpoI is degraded through the destabilization domain (DD). Addition of Shield-1 allows for stabilization of the DD and subsequent I-PpoI protein accumulation in the cytoplasm. Addition of 4-OHT binding to the estrogen receptor (ER) allowing for nuclear translocation and activation of I-PpoI to induce rDNA DSBs. **(B)** Western blot of pS139-H2AX and total histone H3 protein levels in wild-type hTERT-RPE1, hTERT-RPE1 *p53*^-/-^ Cas9 cells, and hTERT-RPE1 *p53*^-/-^ Cas9 cells expressing DD-HA-ER-I-PpoI. **(C)** Quantification of pS139-H2AX/total H3 signal intensity of western blots shown in (A). N=3. **(D)** Immunofluorescet microscopy of hTERT-RPE1 *p53*^-/-^ Cas9 cells or cells expressing DD-HA-ER-I-PpoI treated with DMSO or Shield-1+4-OHT for 6 hr to induce nucleolar segregation or transfected with a plasmid containing *LacZ* or 5’-ETS rDNA targeting sgRNAs. Endogenous TCOF1 and γH2AX are stained in green and red, respectively. DAPI is shown in blue. Scale bar = 10 μm. Individual channels for TCOF1 and γH2AX are shown on the right. **(E)** Cell survival and LD20 determination of hTERT-RPE1 *p53*^-/-^ Cas9 cells expressing DD-HA-ER-I-PpoI. Cells were passaged every 3 days and treated with Shield-1+4-OHT 24 hr after plating for 1 hr at 37°C to include rDNA damage and nucleolar segregation. After 1 hr, cells were washed with PBS and media was replaced. Total number of viable cells were measured by counting as compared to DMSO control treated cells. N=3 biological replicates for 50 nM to 750 nM concentrations, N=2 for 1-2 μm concentrations.

**Figure S2.**
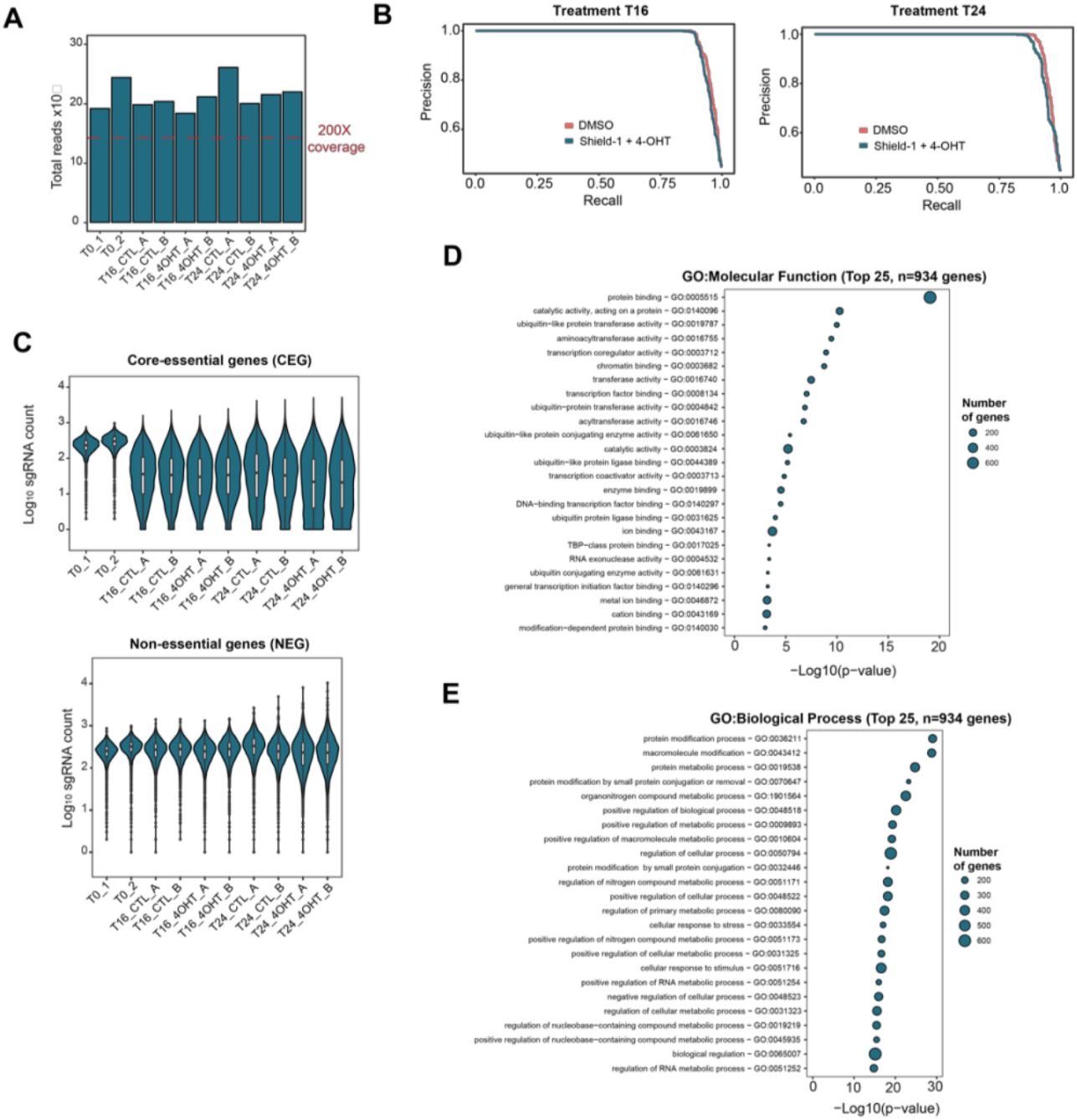
rDNA CRISPR-Cas9 screen quality control and Gene Ontology enrichment. **(A)** Total sgRNA coverage at each timepoint of the rDNA damage CRISPR-Cas9 screen. A minimum of 200X coverage was obtained across all timepoints and samples. **(B)** Precision recall curves at T16 and T24 timepoints of the rDNA damage screen. Curves corresponding to DMSO or Shield-1+4-OHT treated cells are indicated in orange and blue, respectively. **(C)** Total sgRNA counts for core/essential (CEG, left) and non-essential genes (NEG, right) at each timepoint of the rDNA damage screen. **(D)** Top 25 GO:Molecular Function terms of the n=934 genes predicted to be essential for rDNA repair with NormZ scores ≤1.5 as determined by DrugZ analysis. **(E)** Top 25 GO:Biological Process terms of the n=934 genes predicted to be essential for rDNA repair with NormZ scores ≤1.5 as determined by DrugZ analysis.

**Figure S3.**
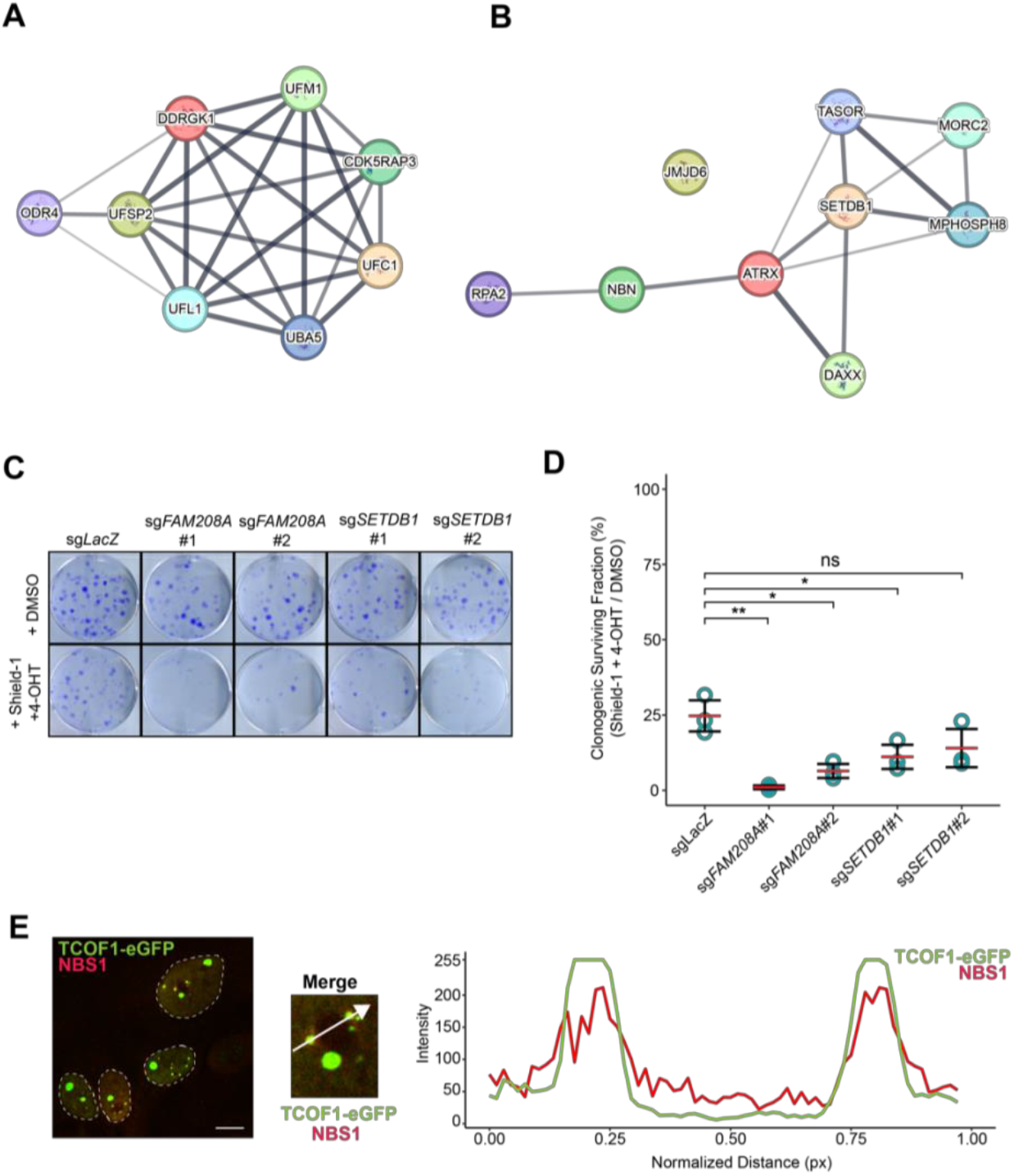
rDNA CRISPR-Cas9 screen STRING maps and HUSH complex validation. **(A)** STRING map of genes known to regulate rDNA as shown in Figure 1D. Edges between proteins represent protein-protein associations such as shared function. Proteins with higher confidence associations are shown in thicker and darker lines. **(B)** STRING map of genes involved in the UFMylation pathway as shown in Figure 1E. Edges between proteins represent protein-protein associations such as shared function. Proteins with higher confidence associations are shown in thicker and darker lines. **(C)** Clonogenic survival assay of hTERT-RPE1 *p53*^-/-^cells expressing DD-HA-ER-I-PpoI transduced with sgRNAs targeting *FAM208A* (encoding TASOR) and *SETDB1* of the HUSH complex. Cells were treated with DMSO or Shield-1+4-OHT every 3 days and allowed to grow for a total of 10-14 days before staining with crystal violet. **(D)** Quantification of the clonogenic surviving fraction shown in (C). Red lines indicate the mean, ±SD is shown (N=3). ns = no significance, * p<0.05, ** p<0.01, unpaired two-sided Student’s t-test compared to sg*LacZ*. **(E)** Zoomed inlet of nucleolar cap as marked with dashed boxes from Figure 4A. White arrow represents the line scan path of relative signal intensity for NBS1 (red) and TCOF1 (green).

**Figure S4.**
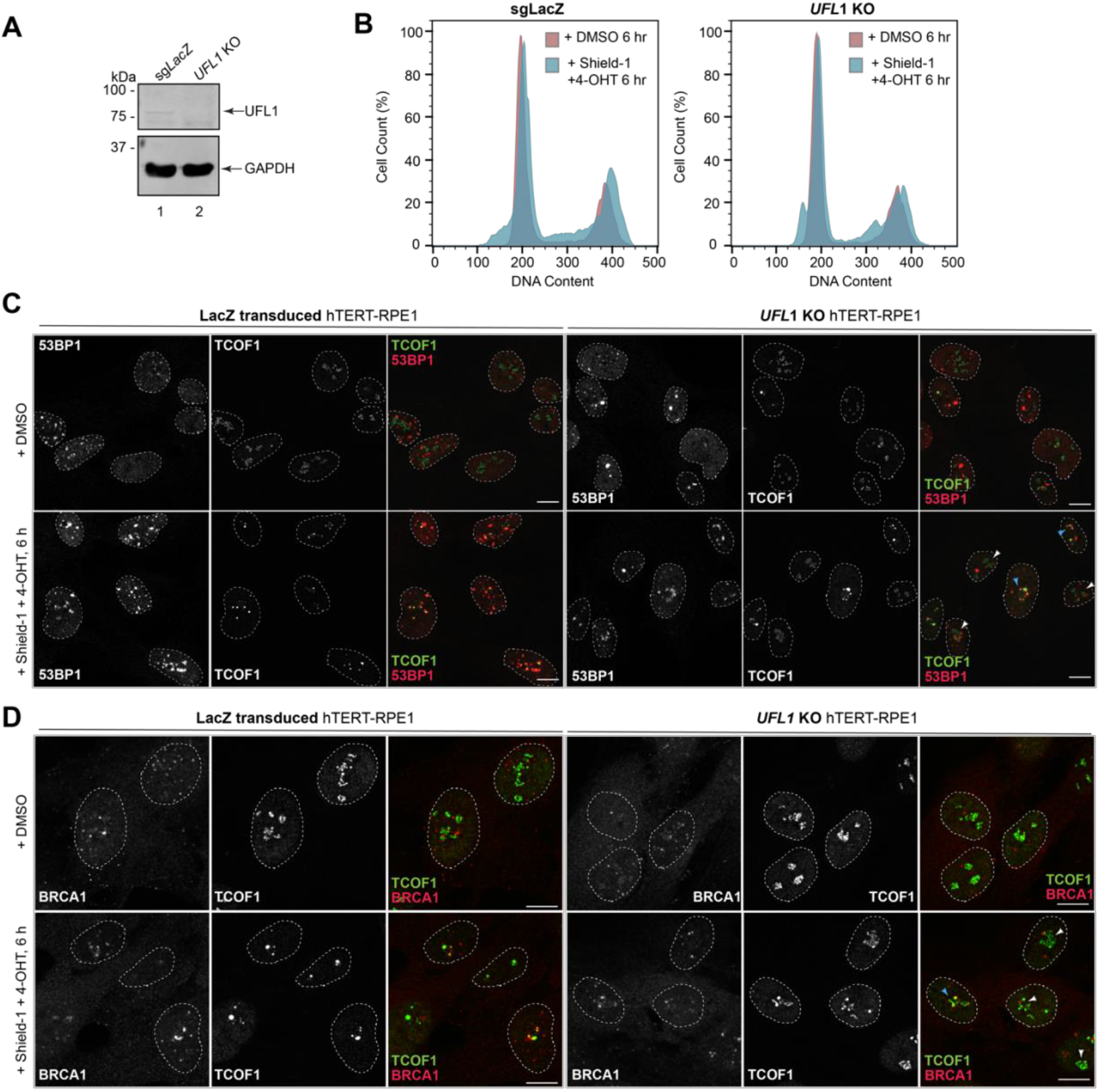
Generation of and characterization of UFL1 KO hTERT-RPE1 cells. **(A)** Western blot of UFL1 and GAPDH protein levels in wild-type hTERT-RPE1 *p53*^-/-^ Cas9 cells expressing DD-HA-ER-I-PpoI transduced with sg*LacZ* or clonal *UFL1* KO cells. **(B)** Cell cycle analysis of wild-type hTERT-RPE1 *p53*^-/-^ Cas9 cells expressing DD-HA-ER-I-PpoI transduced with sg*LacZ* or clonal *UFL1* KO cells. Cells were treated with DMSO (red outlines) or Shield-1 + 4-OHT for 6 hr (blue outlines) before fixation in 70% ethanol. Fixed cells were treated with RNaseA and stained with propidium iodide. Representative cell cycle plots from N=3 experiments are shown. **(C)** Immunofluorescent microscopy analysis of *LacZ* transduced hTERT-RPE1 *p53*^-/-^ Cas9 cells or clonal *UFL1* KO cells expressing DD-HA-ER-I-PpoI treated with DMSO or Shield-1+4-OHT for 6 hr to induce nucleolar segregation. Endogenous TCOF1 and 53BP1 are stained in green and red, respectively. Scale bar = 10 μm. Single z-stack images are shown. White arrowheads: non-segregated nucleoli; blue arrowheads: fully segregated nucleoli/nucleolar cap formation. **(D)** Immunofluorescence micrographs of LacZ transduced hTERT-RPE1 *p53*^-/-^ Cas9 cells or clonal *UFL1* KO cells expressing DD-HA-ER-I-PpoI treated with DMSO or Shield-1+4-OHT for 6 hr to induce nucleolar segregation. Endogenous TCOF1 and BRCA1 are stained in green and red, respectively. Scale bar = 10 μm. Single z-stack images are shown. White arrowheads: non-segregated nucleoli; blue arrowheads: fully segregated nucleoli/nucleolar cap formation.

**Figure S5.**
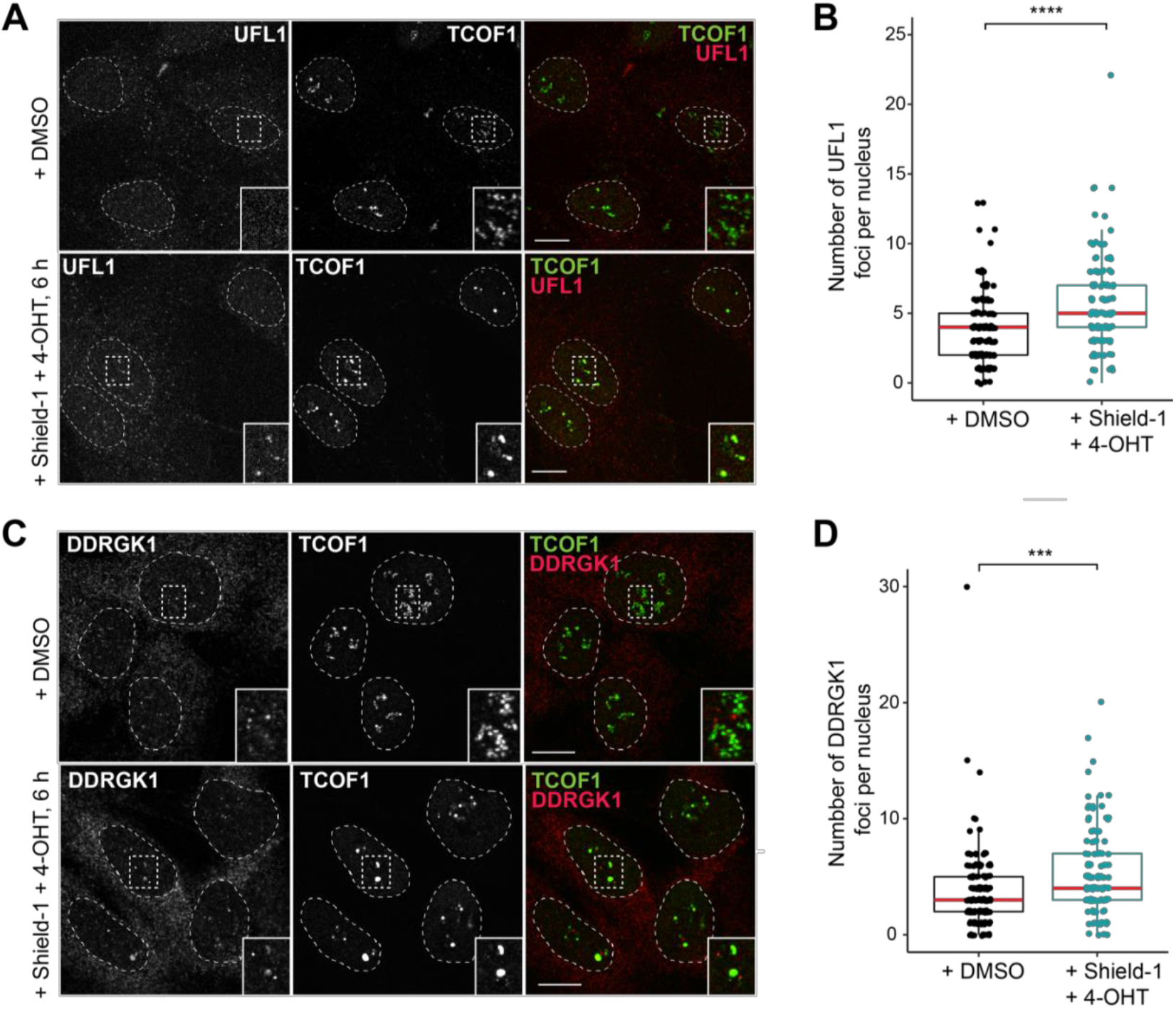
Quantification of nuclear UFL1 and DDRGK1 foci in response to rDNA DSBs. (**A**) Immunofluorescent microscopy analysis of hTERT-RPE1 p53-/-Cas9 cells stably expressing DD-HA-ER-I-PpoI as shown in Figure 5A). Cells were treated with DMSO or Shield-1+4-OHT for 6 hr to induce nucleolar segregation. Endogenous UFL1 and TCOF1 staining are shown individually, with the merged image depicting UFL1 in red and TCOF1 in green, respectively. Zoomed inlets of individual channels are shown. Single z-stack images are shown, scale bar = 10 μm. (**B**) Quantification of total nuclear UFL1 foci from images shown in (A). Red lines indicate median. **** p<0.0001, paired two-sided Student’s t-test compared to DMSO control. (N=3, >125 cells). (**C**) Immunofluorescent microscopy analysis of hTERT-RPE1 p53-/-Cas9 cells stably expressing DD-HA-ER-I-PpoI as shown in Figure 5A. Cells were treated with DMSO or Shield-1+4-OHT for 6 hr to induce nucleolar segregation. Endogenous DDRGK1 and TCOF1 staining are shown individually, with the merged image depicting DDRGK1 in red and TCOF1 in green, respectively. Zoomed inlets of individual channels are shown. Single z-stack images are shown, scale bar = 10 μm. (**D**) Quantification of total nuclear DDRGK1 foci from images shown in (C). Red lines indicate median. *** p<0.001, paired two-sided Student’s t-test compared to DMSO control. (N=3, >125 cells).

**Figure S6.**
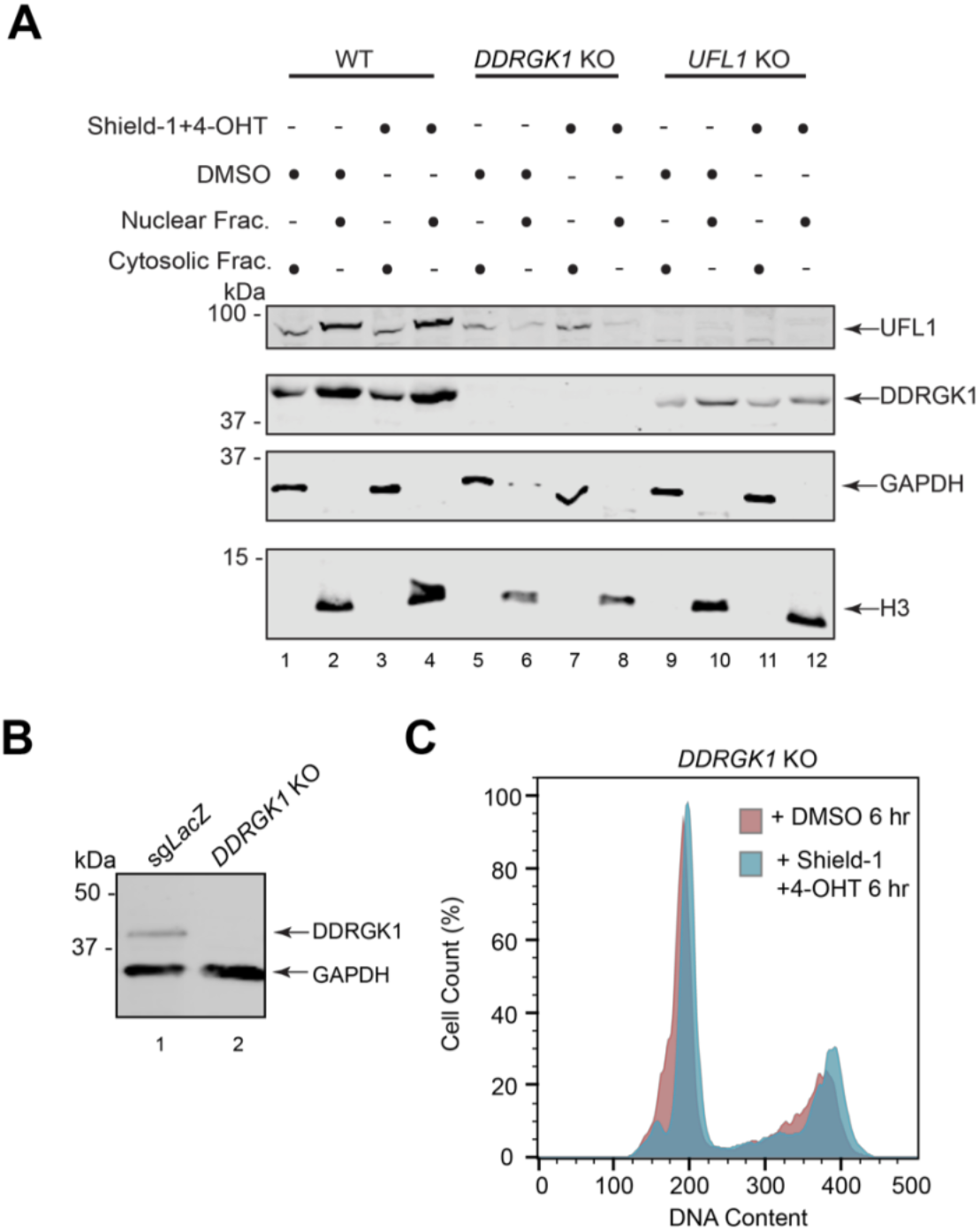
Generation of DDRGK1 KO hTERT-RPE1 cells and characterization of nuclear protein levels of UFL1/DDRGK1 in response to rDNA DSBs. **(A)** Western blot of UFL1 and GAPDH protein levels in wild-type (WT) hTERT-RPE1 *p53*^-/-^ Cas9 cells expressing DD-HA-ER-I-PpoI transduced with sg*LacZ* or clonal *DDRGK1* KO cells. **(B)** Cell cycle analysis of wild-type hTERT-RPE1 *p53*^-/-^ Cas9 cells expressing DD-HA-ER-I-PpoI transduced with sg*LacZ* or clonal *DDRGK1* KO cells. Cells were treated with DMSO (red outlines) or Shield-1 + 4-OHT for 6 hr (blue outlines) before fixation in 70% ethanol. Fixed cells were treated with RNaseA and stained with propidium iodide. Representative cell cycle plots from N=3 experiments are shown. **(C)** Western blot of UFL1, DDRGK1, GAPDH and H3 protein levels in cytosolic and nuclear fractions of wild-type hTERT-RPE1 *p53*^-/-^ Cas9 cells expressing DD-HA-ER-I-PpoI or clonal *UFL1* and *DDRGK1* KO cells. Cells were treated with DMSO or Shield-1+4-OHT to induce rDNA DSBs for 6 hrs at 37°c prior to fractionation.

